# Local and global analysis of macromolecular Atomic Displacement Parameters

**DOI:** 10.1101/2020.06.17.158089

**Authors:** Rafiqa C Masmaliyeva, Kaveh H Babai, Garib N Murshudov

## Abstract

This paper describes the global and local analyses of Atomic Displacement Parameters (ADP) of macromolecules solved and refined using X-ray crystallography method. It is shown that the distribution of ADPs follows the (mixture of) Shifted Inverse Gamma distribution(s). The parameters of the mixture of SIGDs are estimated using Expectation/Maximisation methods. In addition, a method for resolution and individual ADP dependent local analysis of neighbouring atoms has been designed. This method facilitates the detection of the mismodelled atoms and indicates potential identity of heavy metal atoms. It also helps in detecting of disordered and/or wrongly modelled ligands. Both global and local analyses can be used to detect errors in atomic structures thus helping in (re)building, refinement and validation of macromolecular structures. It can also serve as an additional validation tool during data deposition to the PDB.

**Synopsis:** Macromolecular atomic B value distributions have been modelled using a mixture of Shifted Inverse Gamma Distribution. Also, B value and resolution dependent local ADP differences have been applied for validation of heavy atoms and ligands.

## 1. Introduction

The ever-increasing number of macromolecular structures solved by crystallographic and cryoEM methods and deposited to the PDB (Berman et al, 2002; Lawson, 2011) requires statistically robust and automatic tools for refinement (Sheldrick, 2008; Adams et al, 2010; Smart et al, 2011; Murshudov et al, 2011), validation (Read et al, 2011) and deposition (Adams et al, 2019). In general, it is relatively intuitive, although challenging, to design tools for validation of atomic positional parameters as they should comply with the basic structural and chemical properties of macromolecules and there are a number of popular tools designed to do just this (Williams et al. 2018; Joosten, 2012; Vriend, 1990). Designing such tools for ADP validation is less intuitive and, although importance of this problem has been stressed by many authors (Rupp, 2009; Merritt, 2011; 2012), currently there are no widely used tools to check and validate ADPs. One of the potential reasons is that ADPs reflect many such shortcomings of the modelling as crystal deficiencies (e.g. anisotropy, modulation, imperfection of crystals), inaccurate assumptions in data acquisition and processing, modelling problems (modelling the mobility of molecules using individual ADPs is essentially equivalent to the assumption that atoms are oscillating independently around their central position and such oscillation is harmonic and moreover, all unit cells behave exactly same way) and intrinsic mobility of atoms within molecules and molecules within crystals (Kuhs, 2006). There are several papers describing the use the ADP distribution as a validation criterion (Carugo, 1998; 2018; Yang et al, 2016). These works are utilising the fact that, to a certain degree, ADPs represent uncertainty of atomic positions (Schneider et al., 2014; Yang et al., 2016). Masmaliyeva and Murshudov (2019), using a simple fact that B values are proportional to the variances of the distribution of atoms around their central position and the use of Inverse Gamma Distribution as a conjugate prior for data from normal distribution (O’Hagan, 1994), proposed to model the behaviour of ADPs using Shifted Inverse Gamma Distribution (SIGD). They have also demonstrated that there are a number of PDB entries where B values exhibit multimodal distribution. There may be a number of reasons for such behaviour. These include:

1. It is an intrinsic property of molecules within their environment (crystal or multi-domain/multi-subunit structures in cryoEM), where different components (subunits/domains) have different number of neighbours to interact with. In such cases, different subunits/domains may have different levels of mobility, and this can be reflected in the B value distribution. It can be expected that each individual structural unit will behave as a SIGD with different parameters.
2. Some parts of the model (loops, ligands or even domains) may have been placed incorrectly. Essentially, such behaviour indicates that there is very weak or no evidence to support the presence of these parts of the structures, and as such they should be considered with extreme care.

If it is assumed that the noise level on the map is approximately constant over the unit cell, then it can be claimed that local signal to noise ratio depends on the height of the local average electron density and that in turns depends on the local mobility of molecules, therefore it can be expected that: 1) if atoms are placed in wrong positions, then during refinement their B values will increase dramatically to reflect absence of the density, as signal to noise ratio in these regions are close or equal to zero; 2) if two or more domains/subunits have different intermolecular and/or crystal contacts, then they will have different ADPs reflecting their mobility, thus, reducing signal to noise ratio and making interpretation of such regions very difficult. In both cases there will be multiple modes of ADP distribution and correspondingly, signal to noise ratio will be different. It means that, at least for some of the crystal structures, local signal to noise ratio and therefore local resolution will vary over the unit cell, being more or less constant (more precisely having the distribution corresponding to the ADP distribution) over the region covered by the atoms corresponding to the same mode. In this work we model multimodal ADP distributions as a mixture of SIGDs which can potentially be used further to identify mismodelled and/or structurally compact regions. This fact, among a number of other odd behaviours of ADPs, has been shown by Rupp (2009) in his fine textbook on Biomacromolecular Crystallography.

Although modelling of overall ADP distribution is a good technique for identification of suspicious/interesting regions of crystal structures, it does not allow identification of individual mismodelled atoms, residues or ligands. To address this problem, we consider local ADP differences in a given crystal structure. In general, it is reasonable to assume that if two atoms are close to each other in space then their mobility and therefore their ADPs should be similar. It makes sense if we consider molecules including waters as an elastic network; an oscillating atom has almost immediate effect on its surrounding. Moreover, if atoms have been modelled correctly then all factors influencing ADPs of an atom should also influence neighbouring atoms. Therefore, dramatic differences between ADPs of atoms close to each other in 3D space can mainly be due to different occupancies of the atoms and/or different atom identity, *i.e.* heavy atoms may have been modelled as light atoms or *vice versa*.

One of the problems is that the meaning of closeness of two ADP values is not entirely clear. For example, difference between 100Å^2^ and 150Å^2^ can be less significant than difference between 10Å^2^ and 15Å^2^. Moreover, the resolution will also affect the significance of these differences. Therefore, to analyse differences between B values of atoms the resolution as well as the ADPs need to be accounted for. Wang (2018) uses a similar idea to analyse the occupancies of atoms for different elements in crystals. Here this idea is used to calculate differences between ADPs as well as potential adjustment of occupancies to make ADPs of neighbouring atoms similar.

### Organisation of the paper

First, mathematical formulation of the problems of modelling of ADP distribution using mixed SIGDs is described and then formulation for analyses of local differences is given. Then the described methods are applied for re-refined structures from the PDB and the results are analysed.

## Materials and methods

### Global ADP analysis

Multimodal ADP distributions are modelled using the mixture of SIGDs. This distribution has a form:

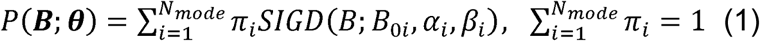

***B*** is a vector of observations, 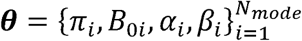 is the vector of parameters, where *π*_*i*_ is the probability of the mode *i. N*_*mode*_ is the number of modes, *SIGD* has the form:

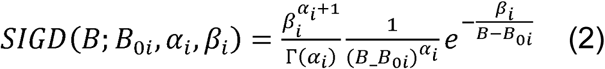

with Γ(*α*_*i*_) as the Gamma function, *B*_*0i*_, *β*_*i*_ and *α*_*i*_ as the shift, scale and shape parameters. The expectation and maximisation algorithm (EM) described by Bishop (2006) is used for the estimation of the parameters of the distribution defined in (1) and (2). Direct application of EM algorithm for the mixture of SIGDs turned out to be unstable. Therefore, the parameters were estimated in four steps: 1) Convert ADP distribution to Peak Height distribution (PHD); 2) use Silverman algorithm (Silverman, 1985) as implemented in *scipy* package to find the number and centroids of the modes; 3) using found number and initial centroids of the modes, fit the mixture of Gaussians into PHD; 4) Starting with the parameters found in the previous steps estimate the parameters of the mixture of SIGDs using EM algorithm (see Appendix A).

### Peak Heights and local ADP analysis

For analysis of relative occupancies of neighbouring atoms peak heights of point atoms with a given resolution and ADPs are considered. In reality the noise level on the amplitudes and phases as well as the weights used in map calculations should also be accounted for. For simplification these factors are ignored. Peak height of a point atom is (Chapman, 1995):

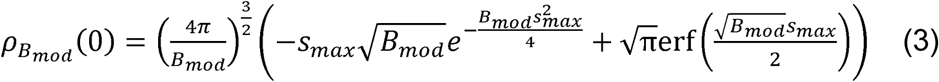

Where *s*_*max*_*=1/d*_*max*_ is the maximum resolution, *B*_*mod*_ is the ADP, *erf* is the error function (for survey of special functions see Abramovitz and Steugun, 1964). Masmaliyeva and Murshudov (2019) use this formula to demonstrate that there is a resolution dependent effect on PHD. If two atoms with ADPs equal to *B*_*1*_ and *B*_*2*_ are considered, then a question can be posed like this: how much the occupancy of the second atom should be changed so that the peak height becomes the same as a fully occupied first atom:

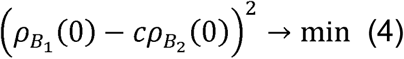

It is solved for *c* to give:

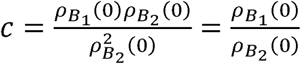

For point atoms putting the expressions from (3) results in:

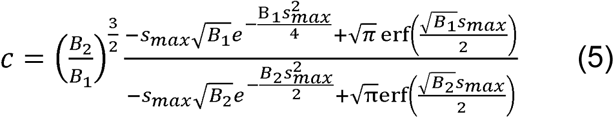

When *s*_*max*_ →∞ this formula converges to:

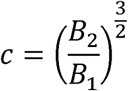

Note that the optimal occupancy value is achieved when the expression in (4) becomes zero, meaning that by changing the occupancies, peak heights at the centre of atoms could be changed arbitrarily. Possible minimum and maximum values of the estimated relative occupancies are *c*=0 and *c*=∞ which are achieved when *B*_*2*_=0 and *B*_*1*_=0 respectively. Obviously, there is no physical meaning for the infinite relative occupancy; it is an artefact of using peak heights at the centre as a guide for atomic identity.

Since the most refinement programs try to fit the total density of atoms into the data, it might be better to use the differences between them to evaluate optimal occupancies. We would like to find the best occupancy for the second atom so that its total density is similar to the first atom:

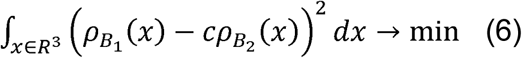

Using Parsewall’s theorem (ignoring constants):

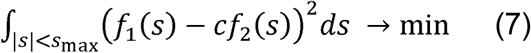

Where *f*_1_(*s*) and *f*_2_(*s*) are scattering factors for the atoms. Solving for *c* gives us:

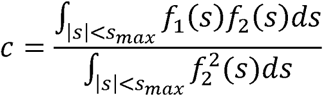

For point atoms this can be written as:

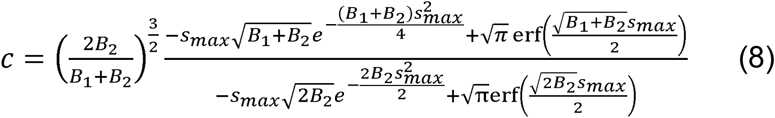

Note that when *s*_*max*_ →∞ then this formula becomes:

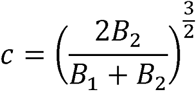

which could be used if the resolution is not known. However, it is recommended to use the resolution, if available, from the PDB entry and set that to the median resolution in all PDB entries (see Table 1) which seems to be around *d*_*max*_=2.1Å. Note that in this case the maximum estimated relative occupancy would be achieved when *B*_*1*_=0 which gives *c*=2^3/2^≈2.83, meaning that, in general, this method will underestimate the occupancy of atoms/ligands/residues. The minimum of (6) is achieved when *B*_*2*_=0 which gives c=0.

**Table 1.** PDB entries rejected from analysis.

| Number of remaining entries | Rejected | Reason for rejection |
| --- | --- | --- |
| 127708 | 12935 | Re-refinement failure* |
| 114773 | 19509 | Outside resolution 1.5–3 Å |
| 95264 | 1491 | High R factor |
| 93773 | 2138 | Viruses |
| 91635 | 795 | No of atoms $n < 500$ |
| 90840 | 13902 | Multimodal |
| **76938** | -- | -- |

\*restrained refinement was applied
Note that Multimodal cases are considered further for modelling using mixture of SIGD

No value of *c* can make the expression in (6) equal to zero unless *B*_*1*_*=B*_*2*_. It means that the only valid explanation of the density is using the correct atoms, which may never be possible. Formulas (5) and/or (8) can be used for quick check of correctness of elements, *e.g.* for Asn, Gln, His side chain orientations. It will only work if the data resolution is sufficiently high and side chains are well defined. In such cases, there will be other atoms around side chains of these residues making H-bonds with them. Therefore, local hydrogen bonding network can be used to correct the orientation of Asn, Gln and His side chains (Chen et al, 2010).

### Data from the PDB

All PDB entries solved by X-ray crystallography, as of November 2019, for which experimental data were available, were downloaded from the PDB and refined using *refmac5* (Kovalevskiy *et al*, 2018) distributed within the CCP4 software suite (Winn *et al*, 2011). The total number of such entries is 127708. All structures were refined with the same software to make sure that all ADPs have been refined consistently using the same software. For this purpose other refinement software could also be used. For further analysis, we used only the models for which the high-resolution diffraction limit is between 1.5Å and 3A□. To avoid dealing with the structures refined with non-crystallographic symmetry constraints, the use of which is not always clear from the PDB, we removed virus structures. Of the remaining models, we were able to refine 90840 automatically. Reasons for refinement failure include (i) the ligand present in the PDB file was not in the CCP4 monomer library (Long *et al*, 2017) at the time of re-refinement - this was the most common case; (ii) absence of experimental data and (iii) space-group inconsistencies between the PDB and data files. We also excluded the cases with R factors > 0.3. Table 1 gives a short summary of the selection of PDB entries. Table 2 lists the examples of the PDB entries used in this work. It should be stressed that the aim of this contribution is not to criticize a particular PDB entry; rather, we would like to highlight the shortcomings of the techniques used at the time of elucidation of these structures and necessity of re-modelling and re-refinement as new technologies become available.

**Table 2.** Summary of the PDB entries used as examples. R and Rfree before, R factors before refinement; R and Rfree after, R factors after refinement.

| PDB code | Case | Resolution (Å) | R before | R after | R<sub>free</sub> before | R<sub>free</sub> after | $\alpha$ | $\beta$ | $B_0$ (Å<sup>2</sup>) |
| --- | --- | --- | --- | --- | --- | --- | --- | --- | --- |
| 5TU8 | 3 modes | 2.33 | 0.210 | 0.201 | 0.240 | 0.250 | 3.49 | 120.58 | 2.31 |
| 4RQZ* | Bimodal | 2.40 | 0.196 | 0.202 | 0.226 | 0.239 | 2.63 | 110.94 | 26 |
| 2WXU | Lighter atom, heavier atom | 1.80 | 0.175 | 0.175 | 0.216 | 0.216 | 4.69 | 95.37 | 3.58 |
| 2ZBL | Heavier atom | 1.60 | 0.152 | 0.135 | 0.182 | 0.160 | 4.85 | 51.89 | 1.32 |
| 5X1O | Wrong ligand | 1.90 | 0.225 | 0.252 | 0.276 | 0.30 | 3.46 | 60.90 | 3.59 |
| 5ORJ | Wrong ligand | 1.99 | 0.20 | 0.209 | 0.221 | 0.244 | 3.96 | 149.25 | 14.01 |
| 6B9B | Wrong ligand | 1.80 | 0.135 | 0.140 | 0.162 | 0.166 | 3.69 | 42.43 | 10.37 |

\*SIGD parameters for multimodal cases are given for unimodal parametrization in this table

## Results and Discussions

The examples below are aimed to demonstrate three aspects of ADPs: 1) modelling of multimodal distributions; 2) identification of mismodelled heavy/light atoms; 3) ligand validation.

### Multimodal ADP distribution

The *α*/*β* plot reported previously (Masmaliyeva and Murshudov, 2019) was recalculated using 76938 structures (Figure 1) with unimodal ADP distributions, the overall picture of the plot is the same as in the previous work.

**Figure 1.**
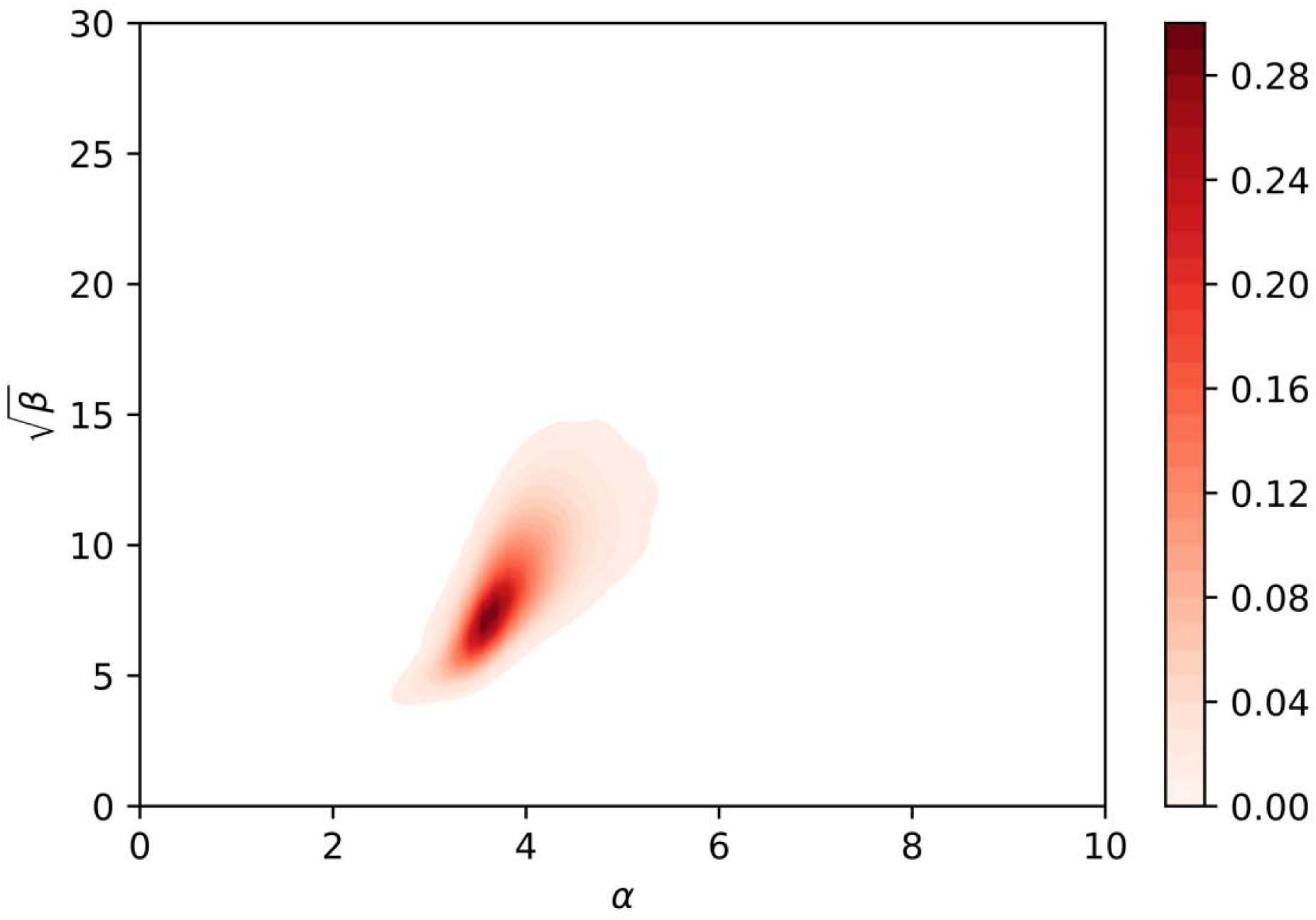
Smoothened of α *vs* β^1/2^ plot for unimodal ADP distributions.

For modes with large centroids *β* values and shift parameters (*B*_*0*_) are high. Also, the ADP distribution corresponding to these modes are more symmetric than the modes with smaller centroids. There are at least two reasons for this: 1) as *β* and centroids become bigger SIGD starts to resemble Gaussian distribution; 2) when ADPs are large then they have large errors. The ADPs correspond to the sum of two random variables – “true” ADP and errors in the estimation. As a result, under a naïve assumption that these two random variables are independent the observed distribution becomes the convolution of SIGD and Gaussian distribution leading to more symmetric distribution.

Estimation of multimodal ADP distributions shows that 13902 out of 90840 cases exhibit multimodality, most of them are bimodal. Because of the reasons given above the second and higher modes are more symmetrical. There are only 266 PDB entries for which ADP distributions show three modes. One such example is 5TU8 (Figure 2). The Figure 2 shows the Gaussian Mixture Model (GMM) for PHD (a), the mixture of SIGDs (b). In the case of 5TU8 the crystal seems to be disordered, part of the crystal does not have any interpretable density, presumably because of very high disorder of the molecules corresponding to this part. The first cluster of ADPs corresponds to the middle part of the molecule, whereas the second and the third clusters correspond to two opposite ends of the molecule where disorder starts. Parameters of the mixture of SIGDs for 5TU8 are given in the Table 3.

**Table 3.**
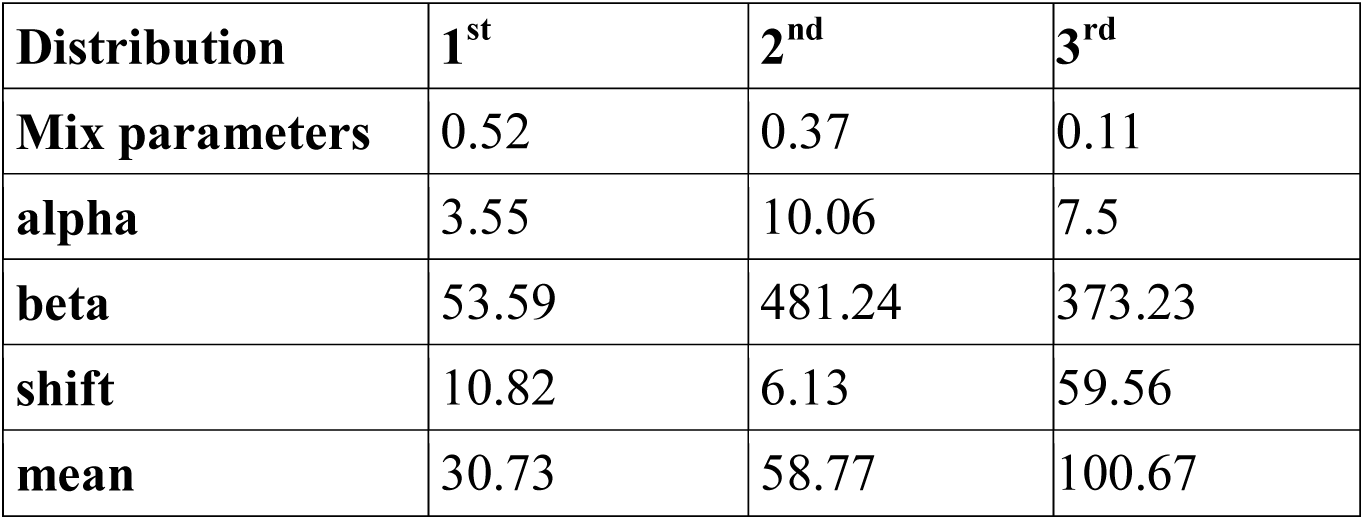
Parameters of the SIGD mixture for 5TU8.

| Distribution | 1<sup>st</sup> | 2<sup>nd</sup> | 3<sup>rd</sup> |
| --- | --- | --- | --- |
| Mix parameters | 0.52 | 0.37 | 0.11 |
| alpha | 3.55 | 10.06 | 7.5 |
| beta | 53.59 | 481.24 | 373.23 |
| shift | 10.82 | 6.13 | 59.56 |
| mean | 30.73 | 58.77 | 100.67 |

**Figure 2.**
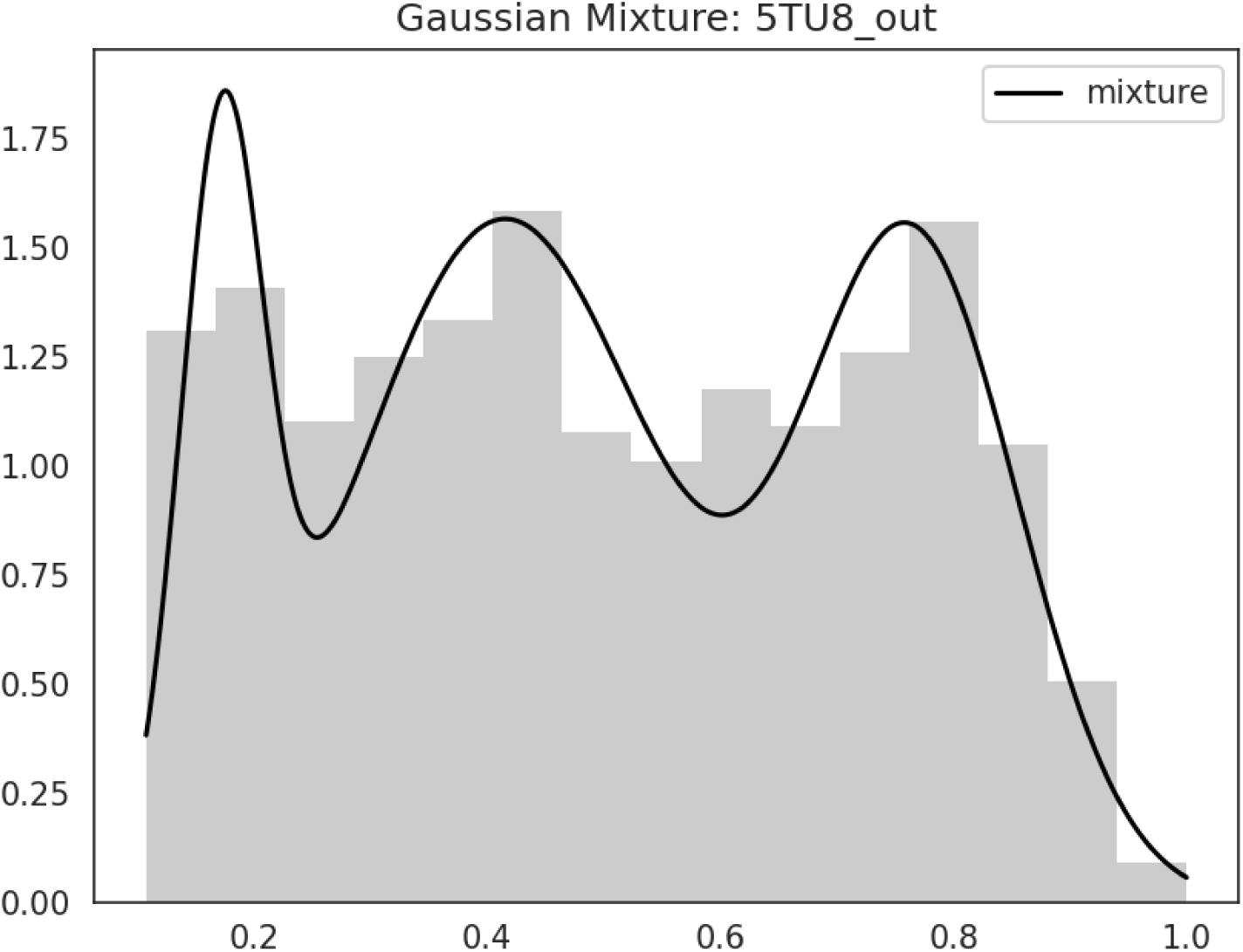
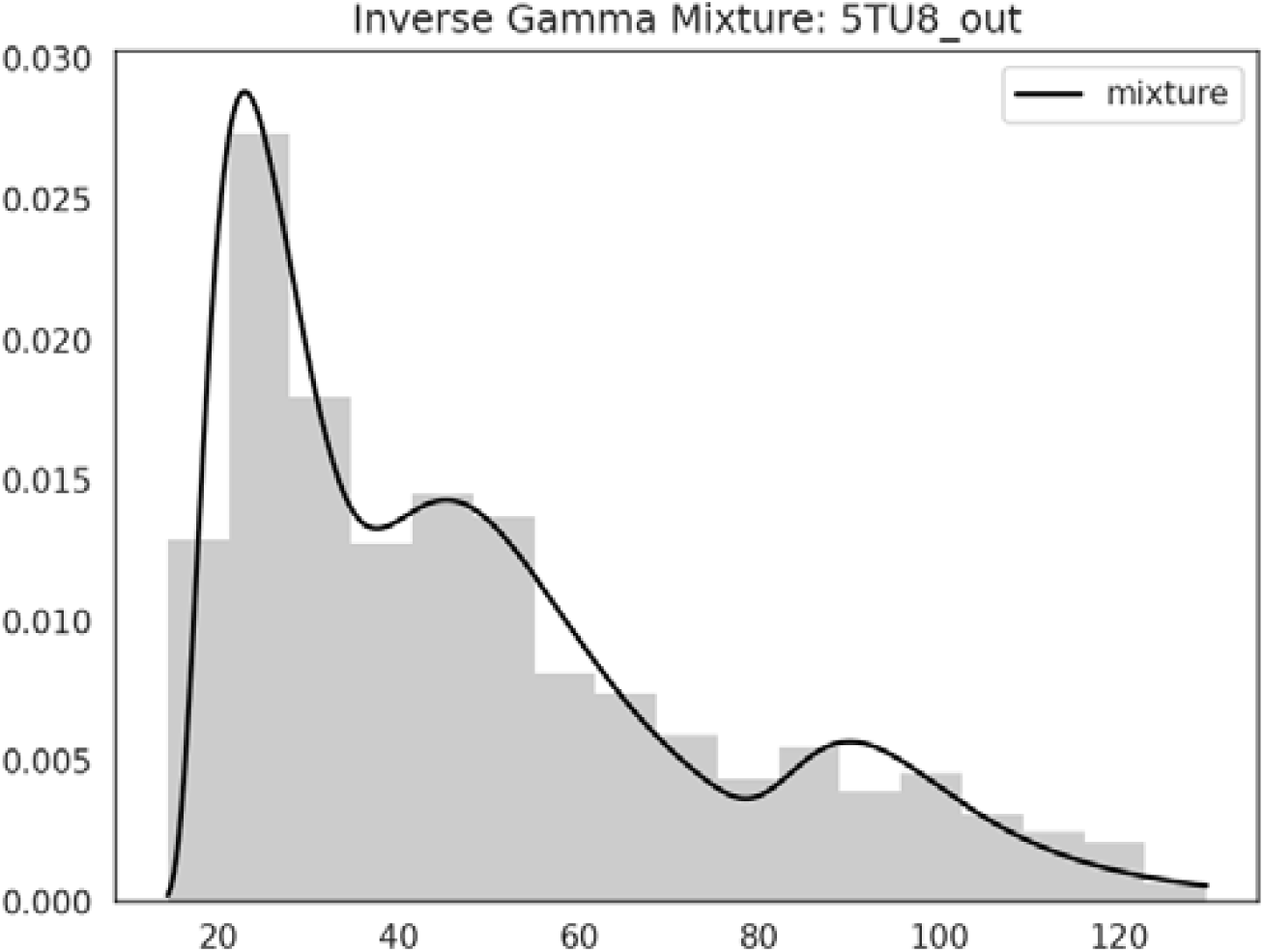
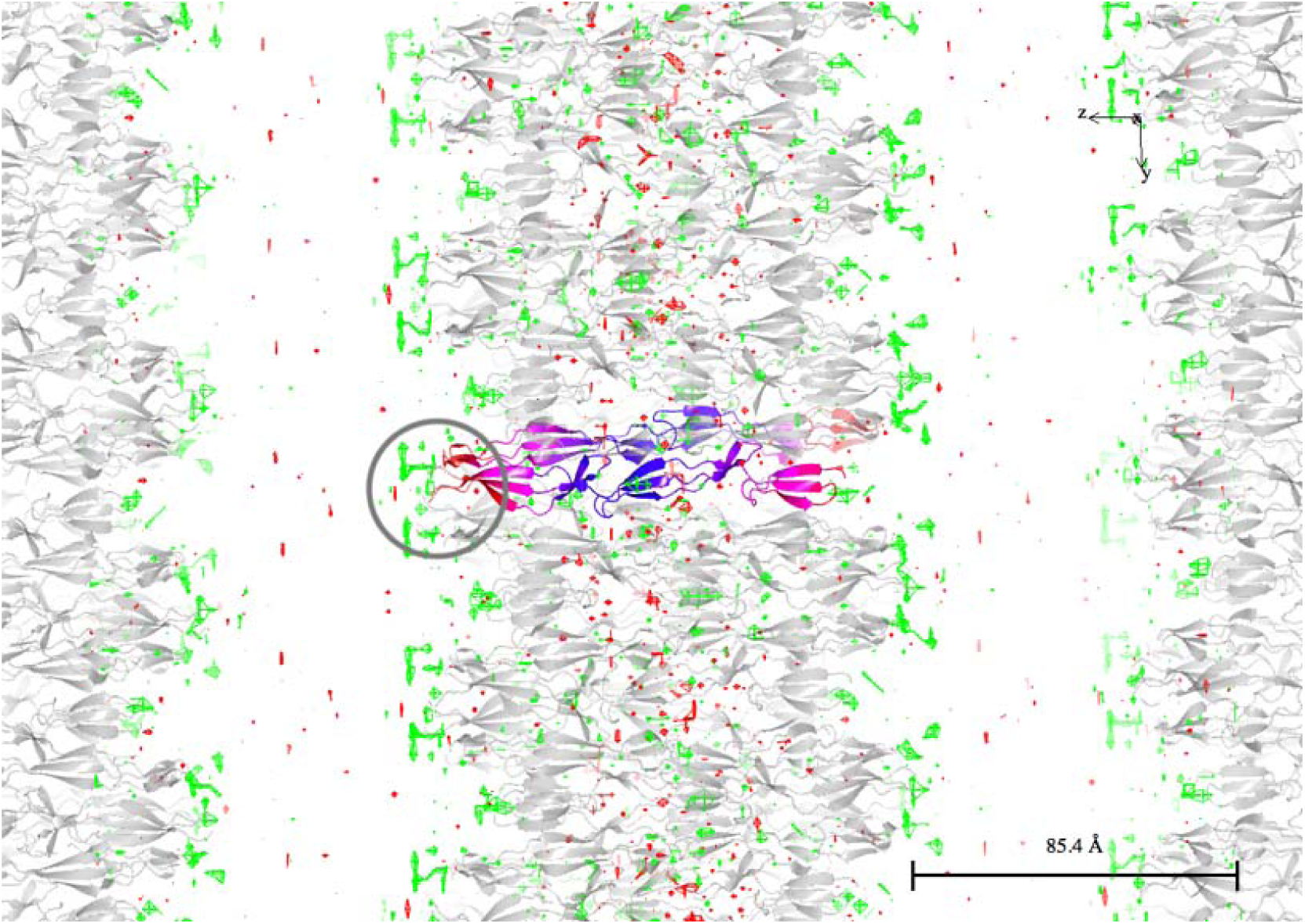
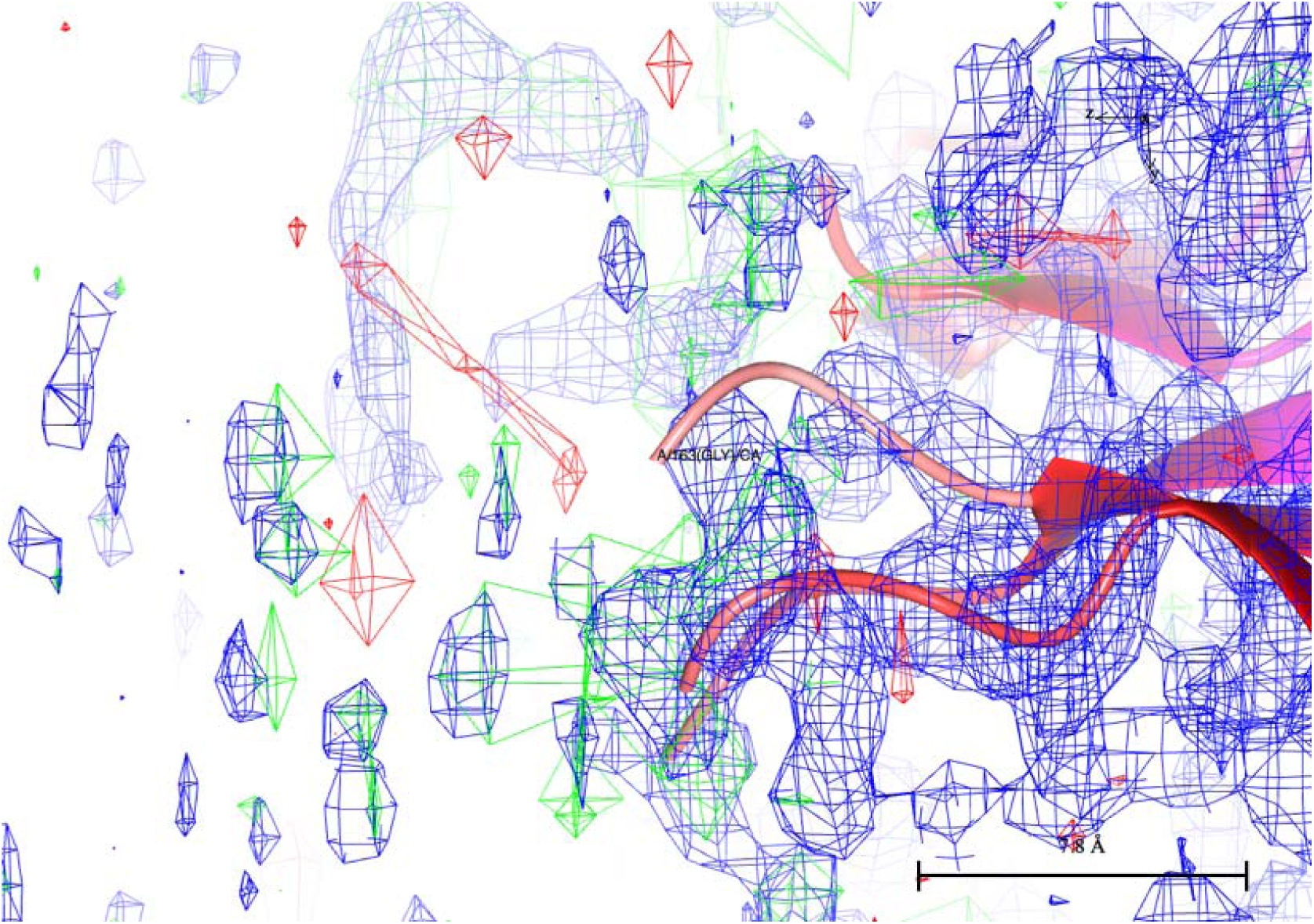
An example of multimodal SIGD with 3 modes – 5TU8. (a) GMM for PHD. (b) The mixture of SIGDs. (c) Continuous crystal for 5TU8 showing disorder. Molecule in the asymmetric unit has been coloured for each cluster – from low to high: blue, magenta, red. (d) The zooming in (marked by an oval in (c)) the end of the molecule shows that there is some positive density there, although it would be a challenge to model them. Figures (c) and (d) were produced using ccp4mg (McNicholas, 2011).

In case 4RQZ entry there are two well defined modes (Figure 3a,b). The molecule has three domains, two of which make contacts with each other and their symmetry mates, these domains are responsible in the crystal formation. The third domain makes contacts only with its symmetry copy (Figure 3c,d). This domain together with its symmetry mate is free to move without further contacts hindering such movements.

**Figure 3.**
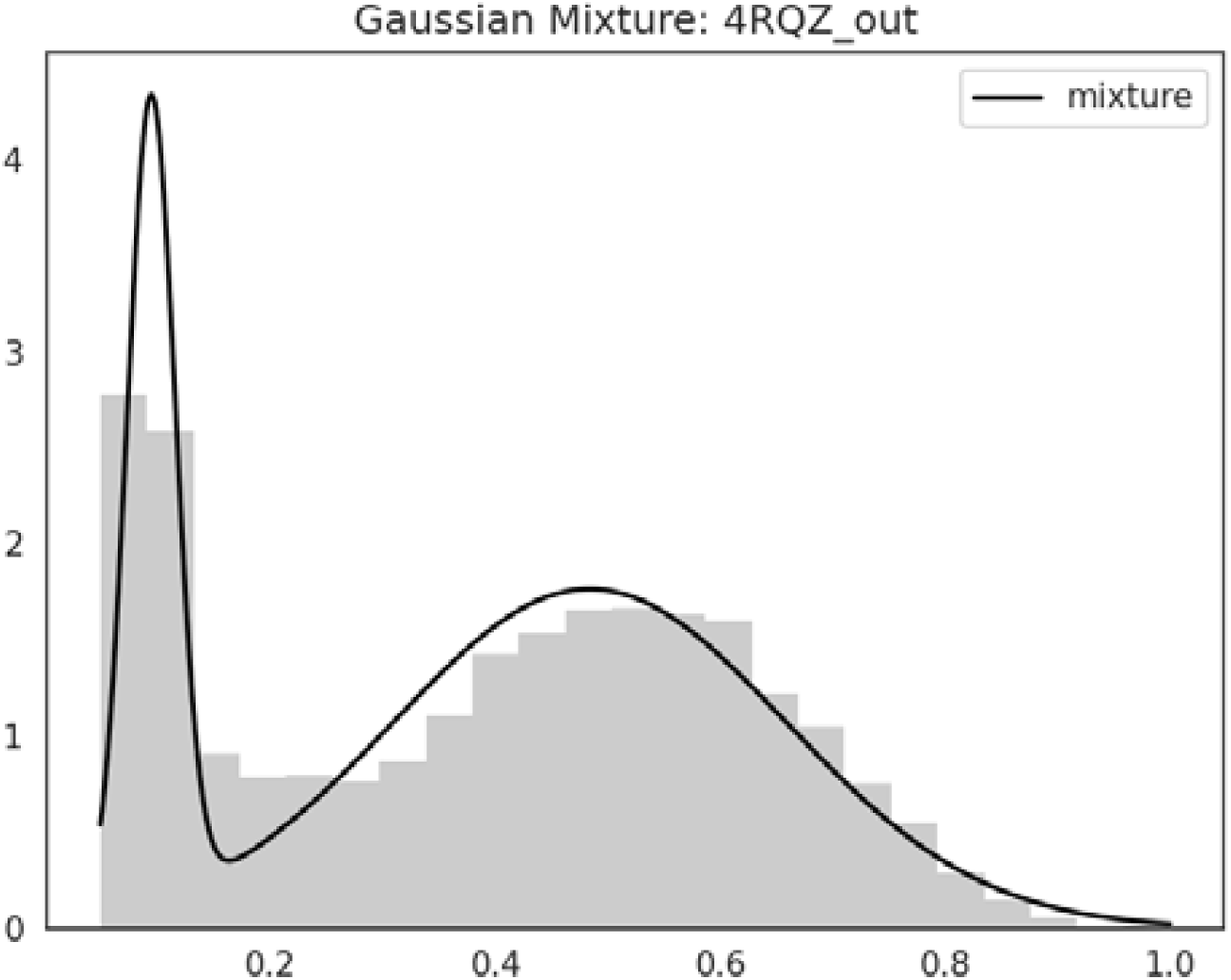
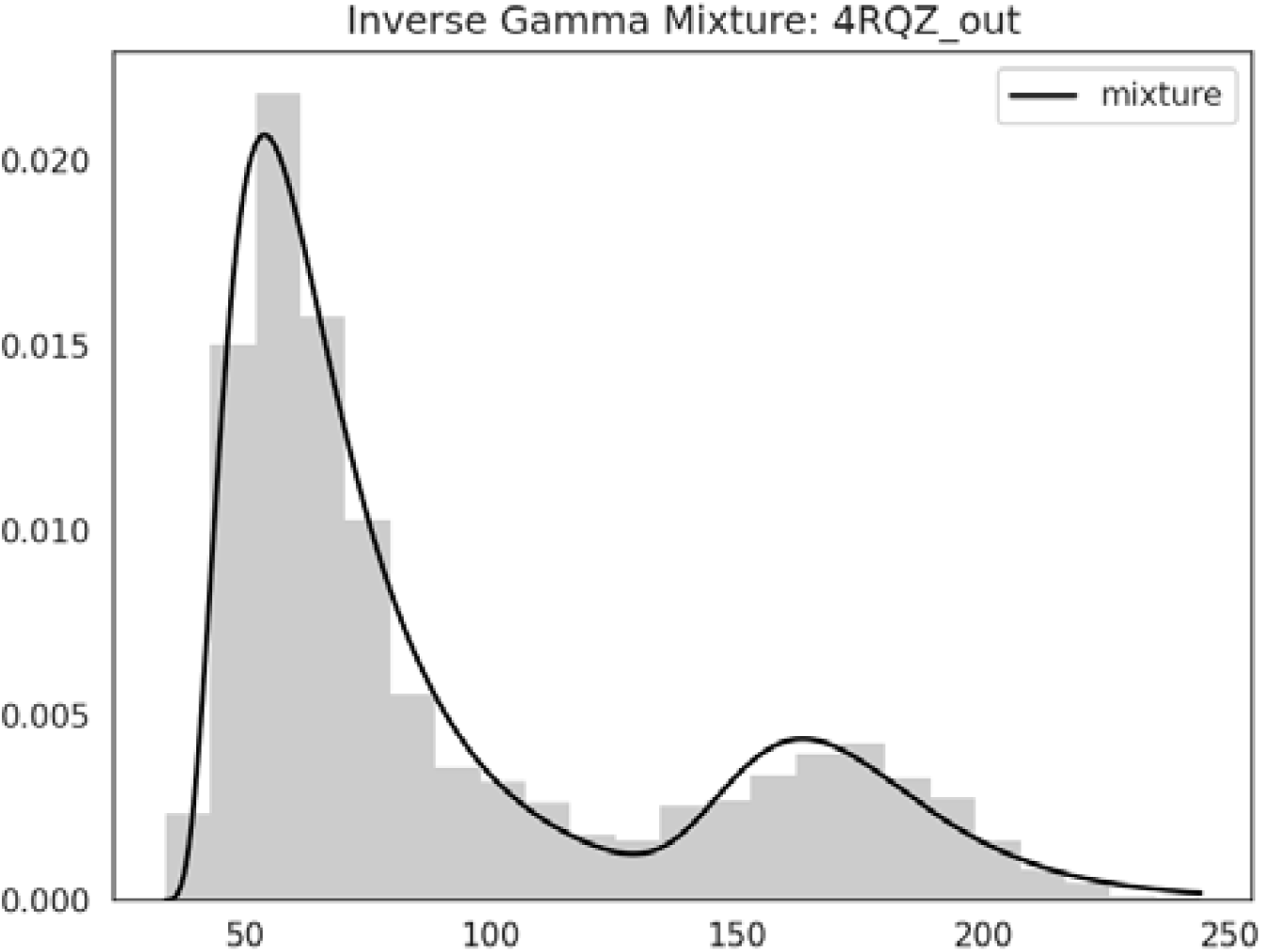
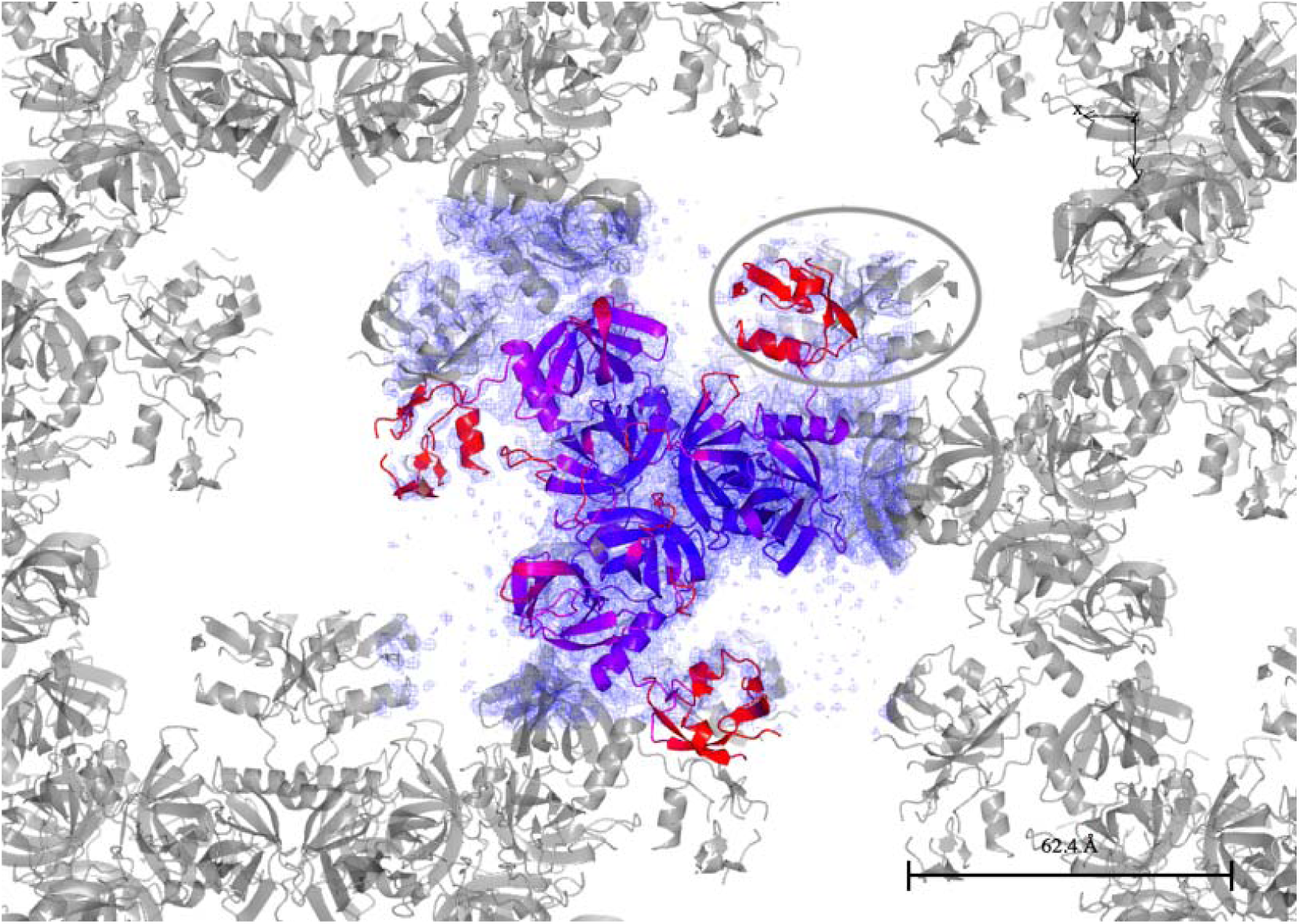
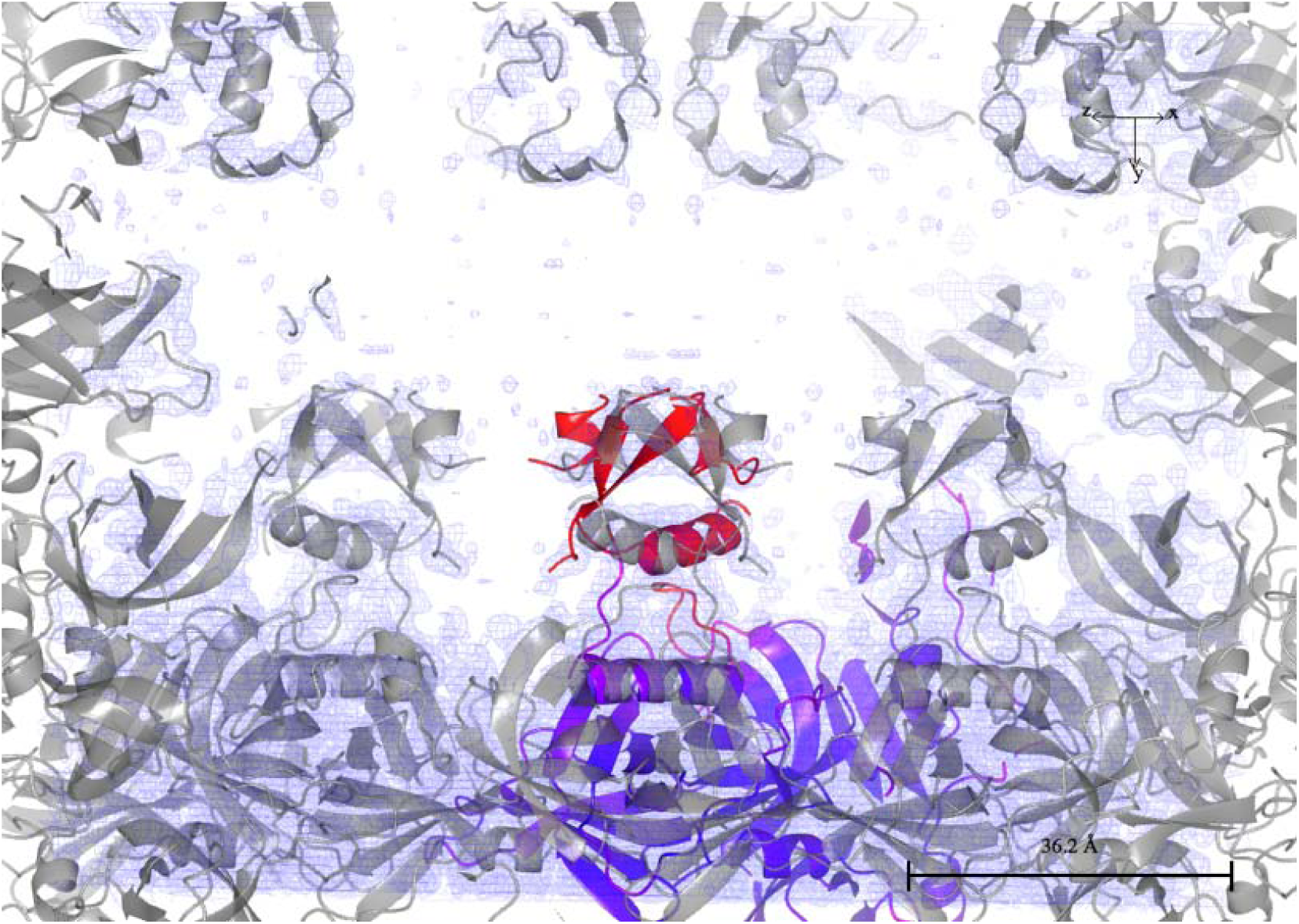
An example of bimodal ADP distribution – 4RQZ. (a) GMM for PHD. (b) The mixture of SIGDs. (c) domains corresponding to the second cluster in the ADP distribution of 4RQZ. (d) crystal contacts of the third domain of 4RQZ, zoomed and rotated version of (c) marked by an oval. Figure (c) and (d) were produced using ccp4mg (McNicholas, 2011)

Parameters of the mixture of B value distributions for 4RQZ are given in the Table 4. As it can be expected the density for the domains corresponding to the second mode is weaker than that for the first two modes (Figure 3).

**Table 4.** Parameters of the SIGD mixture for 4RQZ.

| **Distribution** | **1st** | **2nd** |
| --- | --- | --- |
| **Mix parameters** | 0.79 | 0.21 |
| **alpha** | 3.74 | 10.65 |
| **beta** | 122.74 | 726.67 |
| **shift** | 28.26 | 99.82 |
| **mean** | 72.34 | 174.3 |

**Table 5.** Ligand validation results.

| **PDB** | **5X1O** | **5ORJ** | **6B9B** |
| --- | --- | --- | --- |
| **Resolution (Å)** | 1.9 | 1.99 | 1.8 |
| **Ligand, residue number, chain** | I3P, 201, A | I6P, 407, A | MAL, 807, B |
| **Optimal occupancy (total density)** | 0.121 | 0.162 | 0.414 |
| **Optimal occupancy (PH)** | 0.089 | 0.079 | 0.267 |
| **median(B) of the ligand** | 124.51 | 300.01 | 99.1 |
| **median(B) of the environment** | 11.99 | 52.15 | 37.785 |

### Local ADP analysis

The algorithm described in the Appendix B was applied to all PDB entries considered and more than 1900 entries with data resolution 2Å and better were analysed manually. More than 600 entries were identified as potentially containing heavy atoms and their densities were studied carefully. The electron density corresponding to the atoms marked as light atoms are weaker and in many cases these atoms are exposed to solvent. As a result, in many cases the exposed atoms have high ADPs than surrounding atoms. Residues containing these atoms could have multiple conformation and they might have been subjected to radiation damage. Analysis of radiation damage is outside of the scope of this work.

One example of potentially lighter than surrounding atoms (CD1 of 131 residue ILE of chain A) in the pdb 2WXU is given in Figure 4. The calculated optimal occupancy is 0.63. The ADP of this atom is 36.94 whereas median of the ADPs of surrounding atoms is 19.83. Figure 4b shows that this residue has been modelled in a wrong rotamer. After correction using the program coot (Emsley et al, 2010) for the rotamer and refinement (Figure 4b) B value of the atom is 31.22 and the estimated occupancy is 0.69. There are still some positive and negative densities around this residue indicating that there might be multiple conformations. However existing data quality does not warrant modelling multiple conformation accurately.

**Figure 4.**
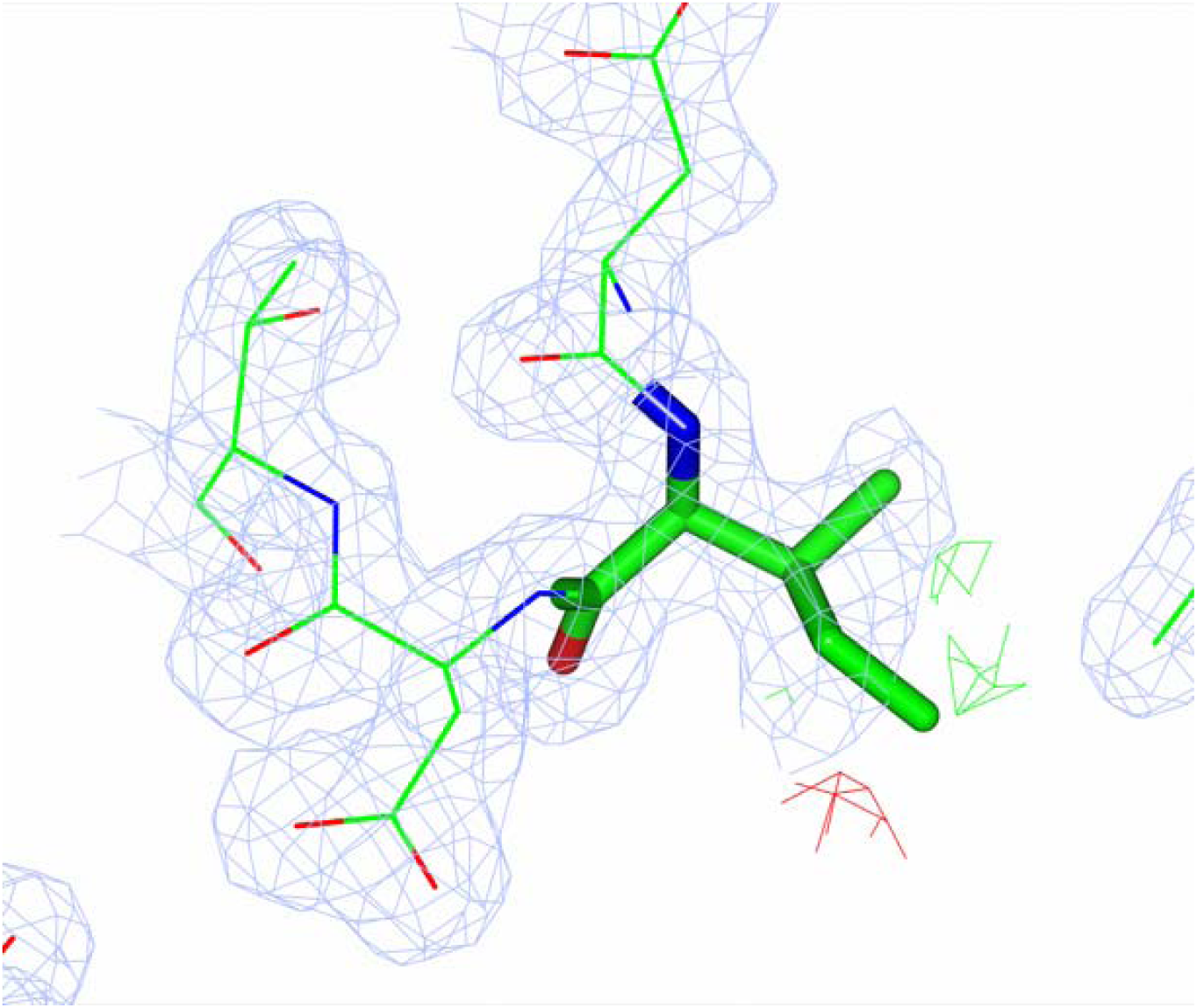
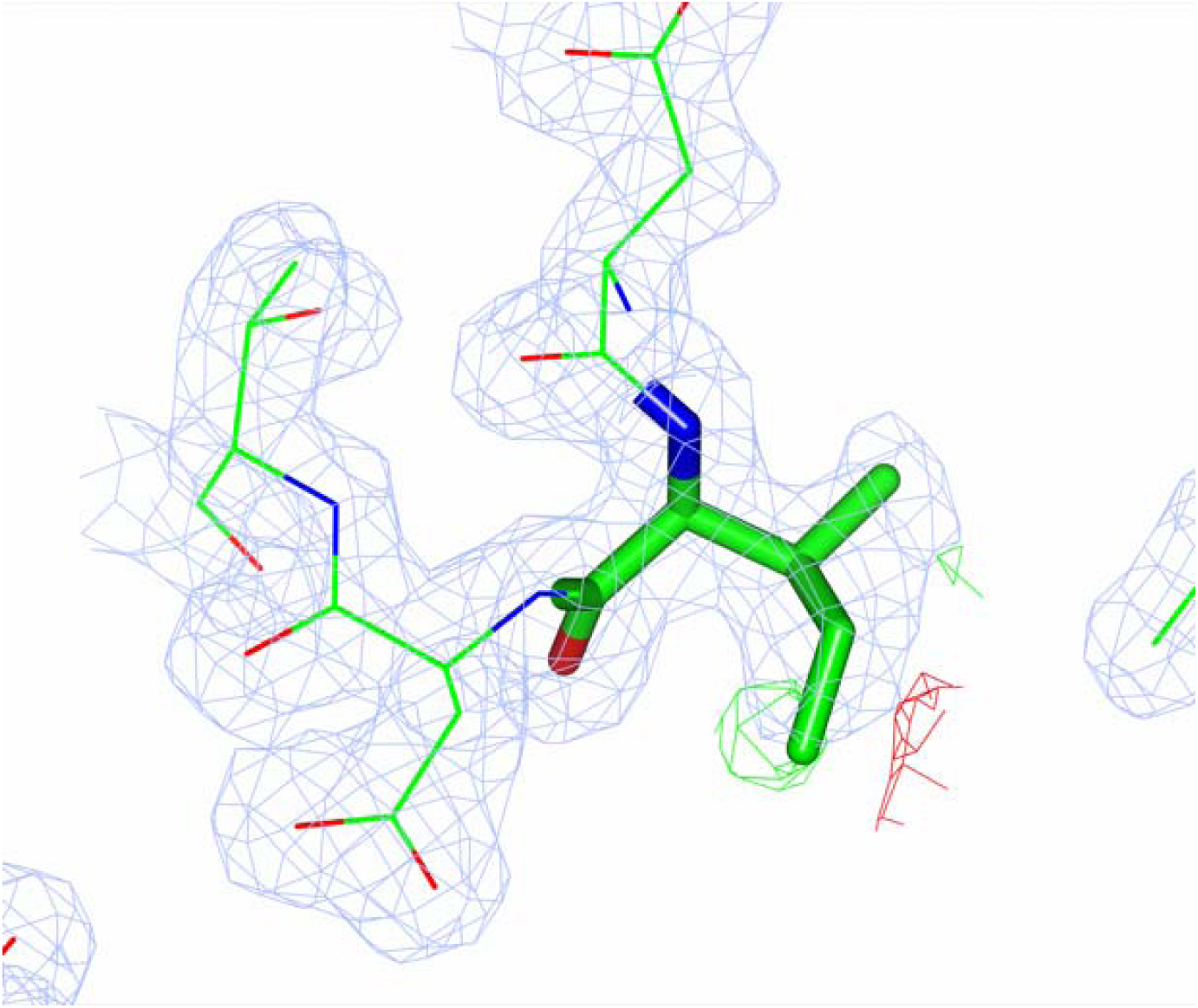
Potential lighter than surrounding atoms - atom CD1 of 131^st^ residue ILE of chain A of 2WXU pdb. (a) Rotamer of ILE as in the PDB file. (b) Rotamer of ILE of after rebuilding. These figures were produced using ccp4mg (McNicholas, 2011)

It is likely that some metals are modelled as water by automatic water picking software as they usually do not analyse the interaction with the surrounding atoms and the height of the electron density. They usually look for the existence of the difference density. The several such cases have been identified in the examined pdb entries. Figure 5 illustrates one such case. In the case of 2ZBL, water molecule of residue 515 chain F had six coordinating atoms forming almost perfect octahedron. The ADP of this atom was 7.22 with the median of surrounding atoms’ ADPs 14.8. It is one of the indicators that it can be a metal atom. The relative occupancy of this atom as calculated using the formula (6) is 1.37. Since it was modelled as O with 8 electrons and 1.37*8 = 10.96 which suggests that this atom can be Na or Mg. Inspection of the crystallization condition showed MgCl_2_ was used in the buffer. It is an indication that Mg is more likely than Na. Analysis of the distances between this atom and surrounding atoms shows that they are between 2.09Å and 2.2Å. Distance between Mg and O is around 2.06 and that between Na and O is around 2.35. Taking all these factors together suggests that it is likely that this atom is Mg. After modelling it as Mg and ten cycle refinement the ADP of this atom became 18.54Å^2^ with estimated occupancy equal to 0.85.

**Figure 5.**
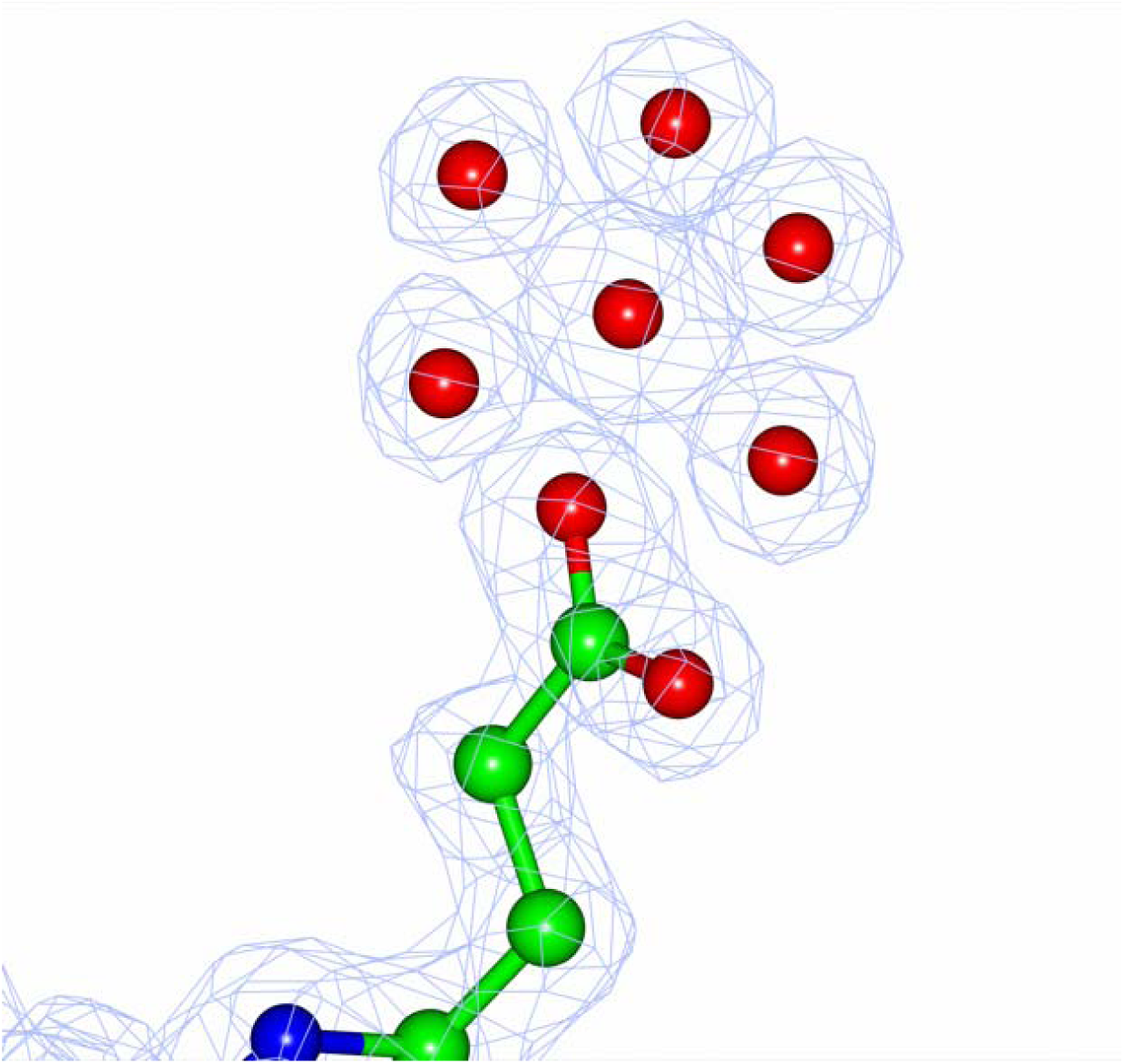
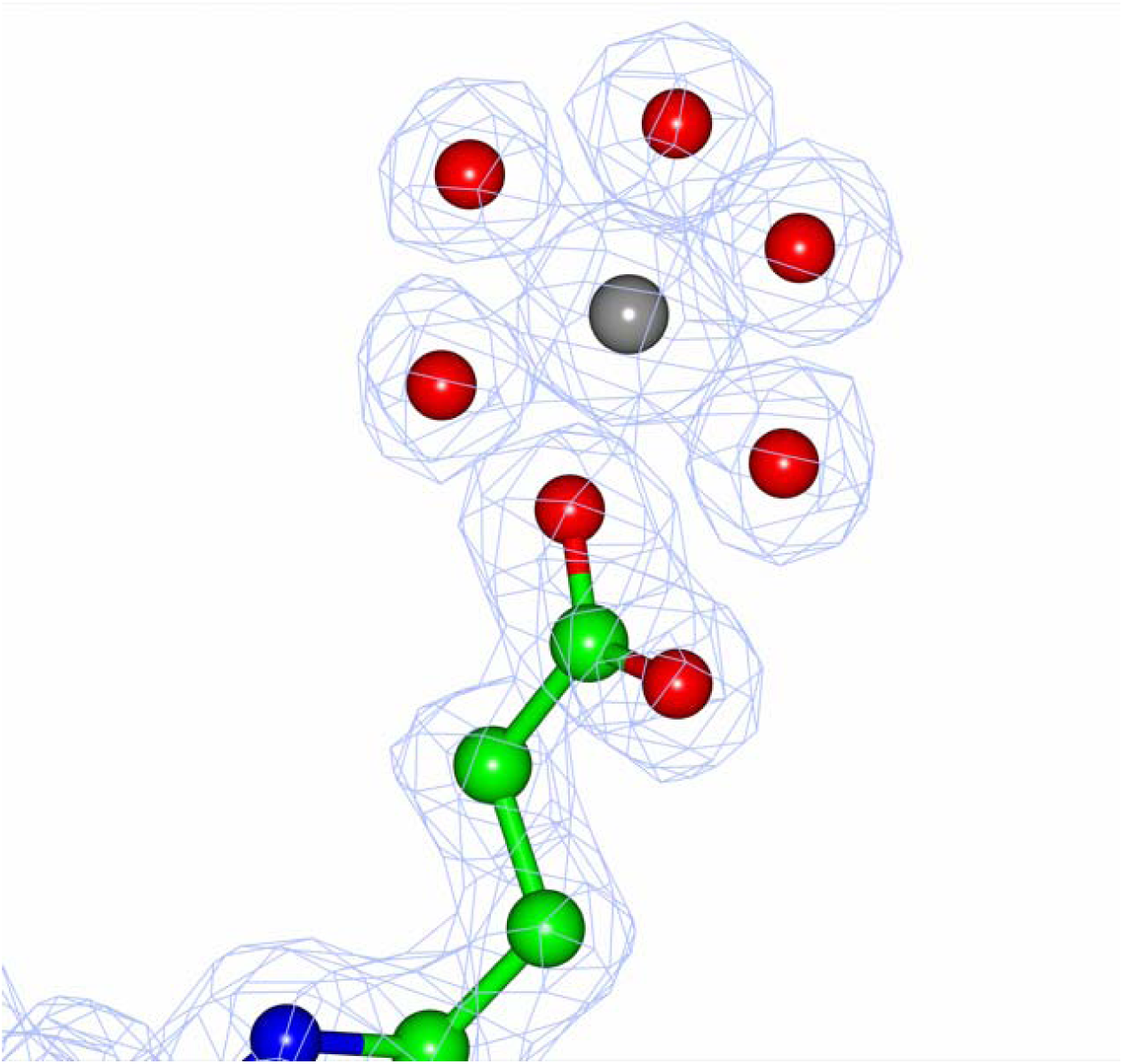
An example of a heavy atom modelled as a water molecule - residue 515 of chain F of 2ZBL, which presumably is Mg. (a) Potentially heavier atom modelled as water. (b) Mg atom in this position after rebuilding and re-refinement. These figures were produced using ccp4mg (McNicholas, 2011)

Many PDB entries contain heavy atoms, most of them seem to have correct parameterisation. However, in some cases these atoms are modelled incorrectly. One such entry is 2WXU, there is Ca atom with the residue number 1377 of chain A, with relative occupancy equal to 1.37. The program marked this as a heavier atom with B value 13.74 and median B values of the neighbours 25.02. Crystallization condition had CdSO_4_. Refining this atom as Cd with half occupancy made the ADP of this atom 18.40 closer to that of surroundings. After re-building using coot (Emsley *et a*l, 2010) and re-refinement, this atom is no longer reported as an outlier. There were still some positive densities around this atom. This atom is close to its symmetry mate, the distance between them is 2.3Å, it is close to the “ideal” distance between Cd and oxygen atoms. It means that when Cd is one of the positions another position is occupied by a water molecule. Surrounding protein atoms also adjust to accommodate Cd/Water switch. Existence of multiple conformations also explains why surrounding atoms have larger ADP than Cd atom. If we consider that Ca atom has 20 electrons then 1.38*20 = 27.6. When there is Cd atom with half occupancy then there are 0.5*48 = 24 electrons. If we add half occupied oxygen atom then we add 4 more electrons giving 24+4 = 28, which is close to 27.6. Obviously, such accurate calculation may not be always possible. To do such calculations the resolution of the data must be sufficiently high and atoms must be well defined. Figure 6 shows this atom together with its symmetry mate and its coordination together with the map.

**Figure 6.**
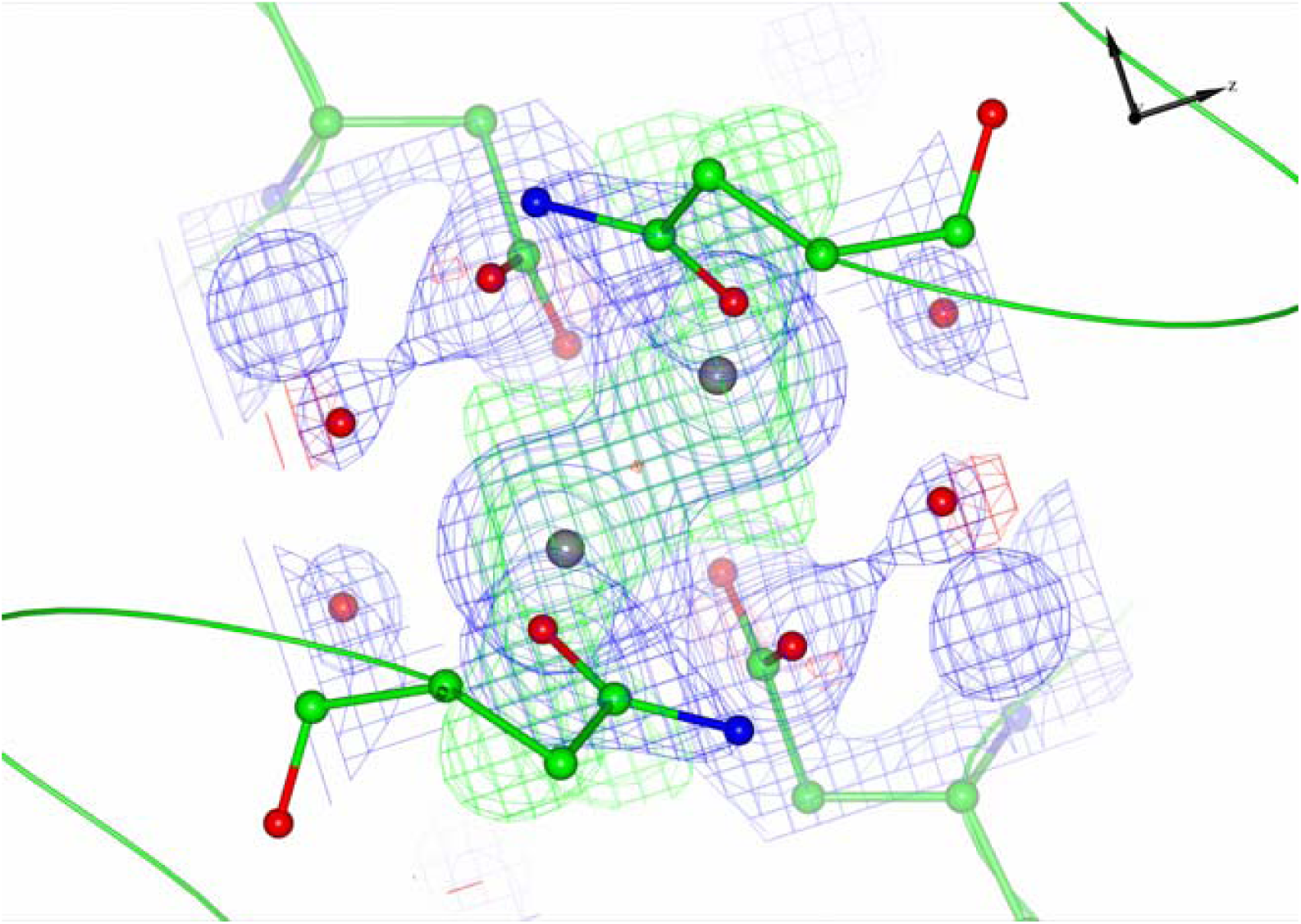
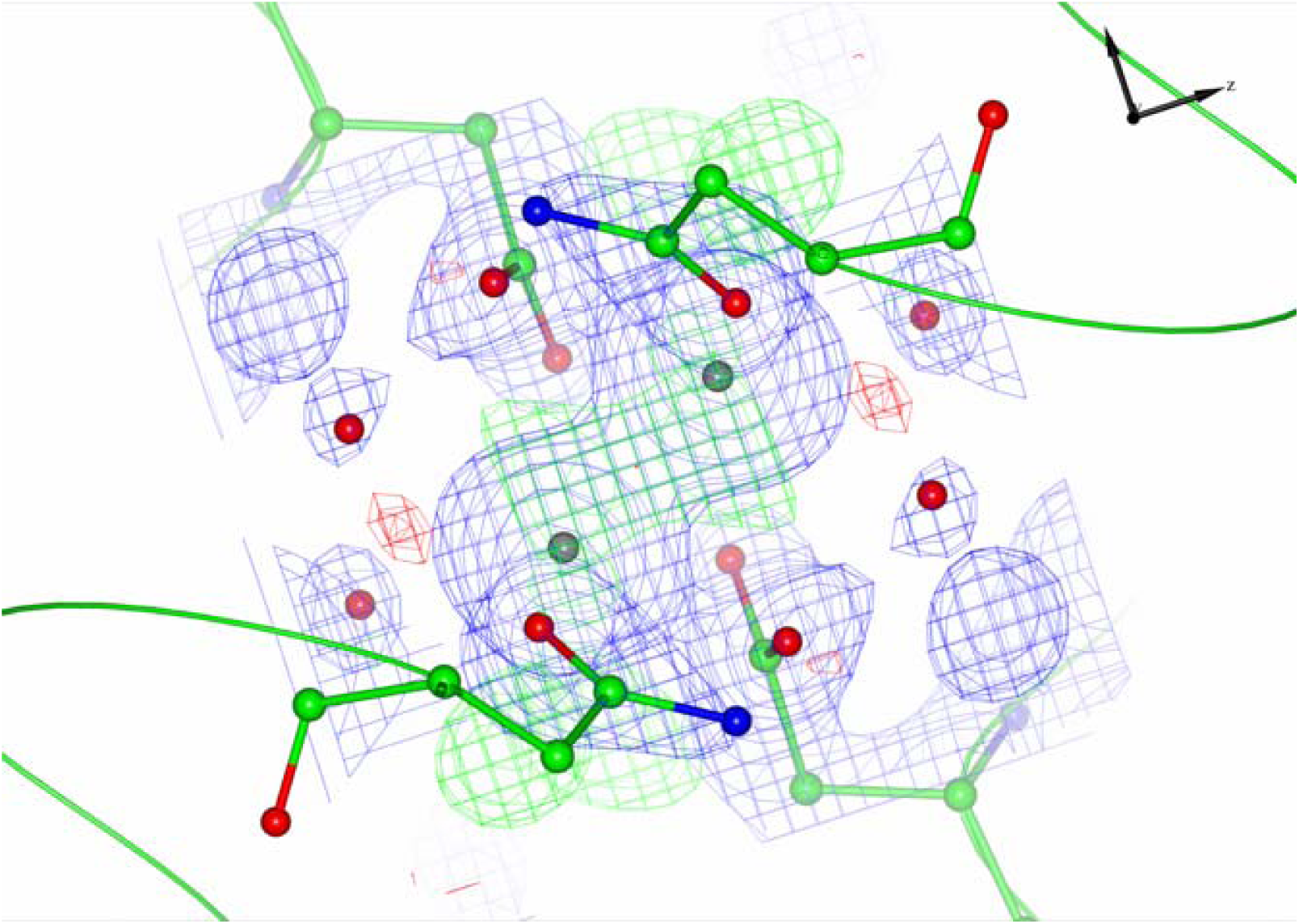
Ca atom of the residue 1377 of chain A in 2WXU detected as heavier than neighbouring atoms. (a) Potentially heavier atom modelled as Ca. (b) Half occupied Cd atom at the same position after rebuilding and re-refinement. In both maps, atoms symmetry mates are coloured using the same colouring scheme. Two-fold crystallographic symmetry axis is perpendicular to the plane and goes through centre of line connecting heavy atoms. These figures were produced using ccp4mg (McNicholas, 2011)

### Application of local ADP analysis to ligand validation

Local analysis was also applied for ligands. In this case all ligand atoms where considered and median ADP of ligand was compared to that of neighbouring atoms. There were number of cases where ligands were marked as having substantially less than full occupancy. There were number of SO_4_ and PO_4_ groups that do not seem to have supporting experimental evidence. We did not consider these cases further. There are number of Zn and other metals with suspected density, since these cases are considered by Touw et. al. (2016) we did not analyse them further. More than ten PDB entries were inspected in detail. Among them three of them were selected for this work. These are 5×1O with ligand I3P –Inositol 1,4,5-trisphosphate, 5ORJ with ligand I6P –Inositol 1,2,3,4,5,6-hexakiphosphate and 6B9B with the ligand MAL – Maltose. Table 6 gives the relative estimated occupancies for these ligands together with the median ADPs of the ligands and surrounding atoms.

#### Case 1: 5×1O

Estimation of occupancy for the ligand I3P of this PDB is 0.11 indicating that this ligand either is not present or present with very low occupancy. Inspection of the electron density showed (Figure 7 a) there is no convincing electron density corresponding to this ligand. After removing this ligand and adding waters where it is necessary the difference map became cleaner (Figure 7b).

**Figure 7.**
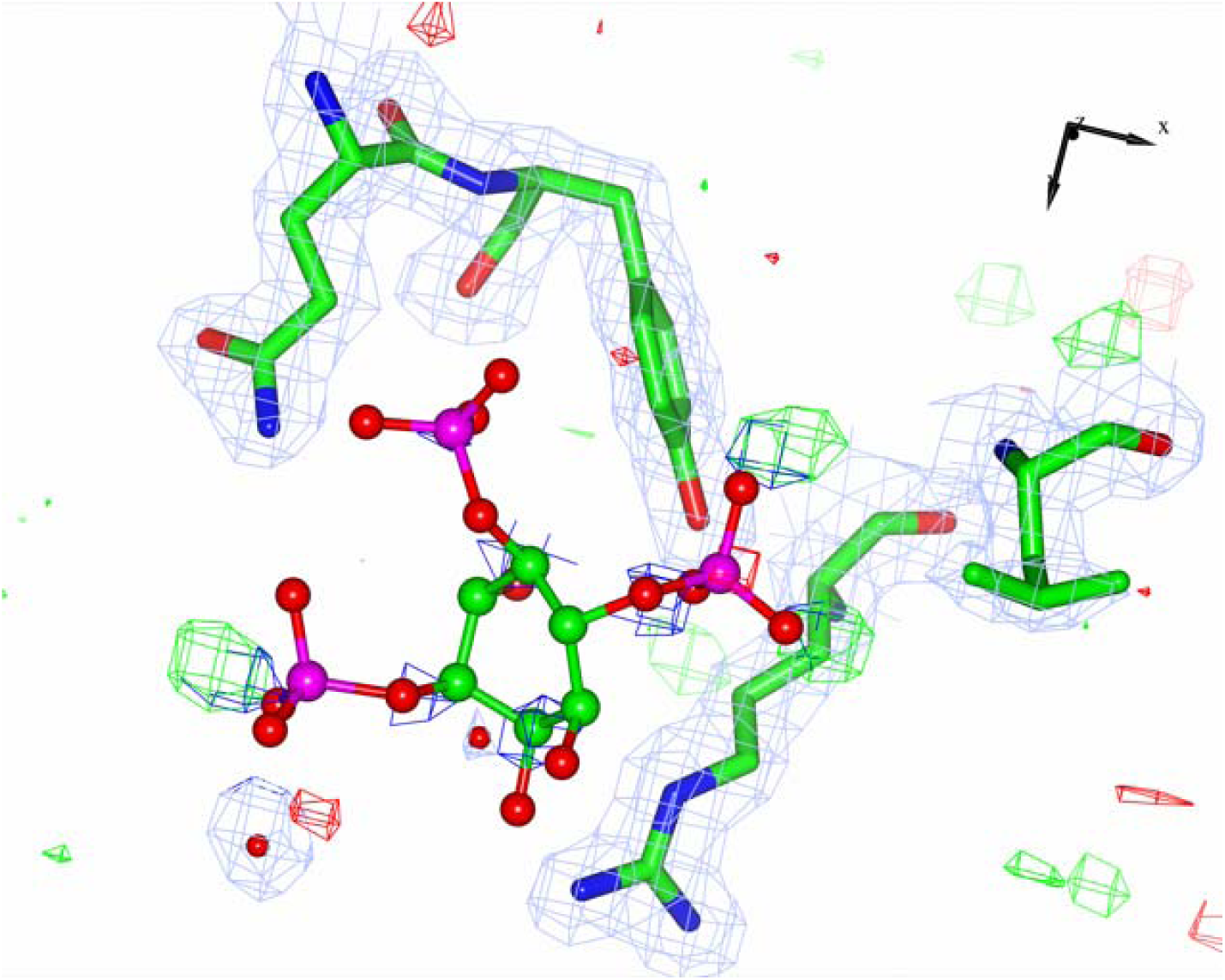
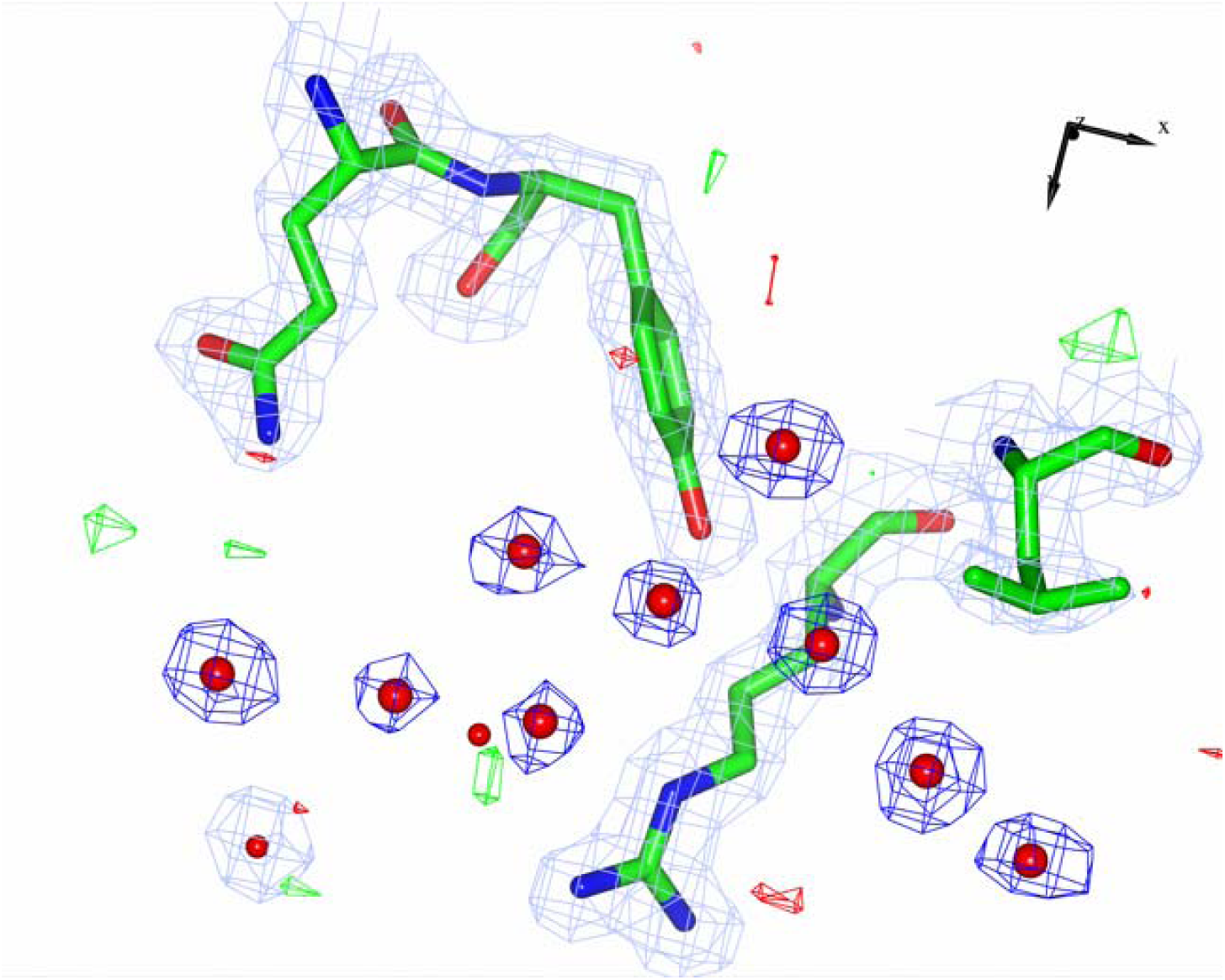
I3P ligand of 5×1O. (a) The ligand before rebuilding and re-refinement. (b) Water molecules placed according to the difference map. These figures were produced using ccp4mg (McNicholas, 2011)

#### Case 2: 5ORJ

Estimated occupancy of the ligand I6P of this ligand is 0.162 which again shows that either it is absent or present with low occupancy (Figure 8a). Its median ADP is 300.01 and that of surrounding atoms is 52.15. Inspection of the density and symmetry related molecules showed that this ligand is on a two-fold-axis resulting of non-bonding repulsions of symmetry related molecules moving them out of the density. After 40 cycles of refinement with half occupancy the median ADP of the ligand became 84.84 with that of the neighbours equal to 51.77 and estimated occupancy became 0.66. The difference density became clean (Figure 8 b), although the density is still weak. It suggests that although half occupied ligand fits better there is still some disorder or mobility of this ligand. It is also a clear demonstration that during model building and refinement crystal symmetries must be accounted for.

**Figure 8.**
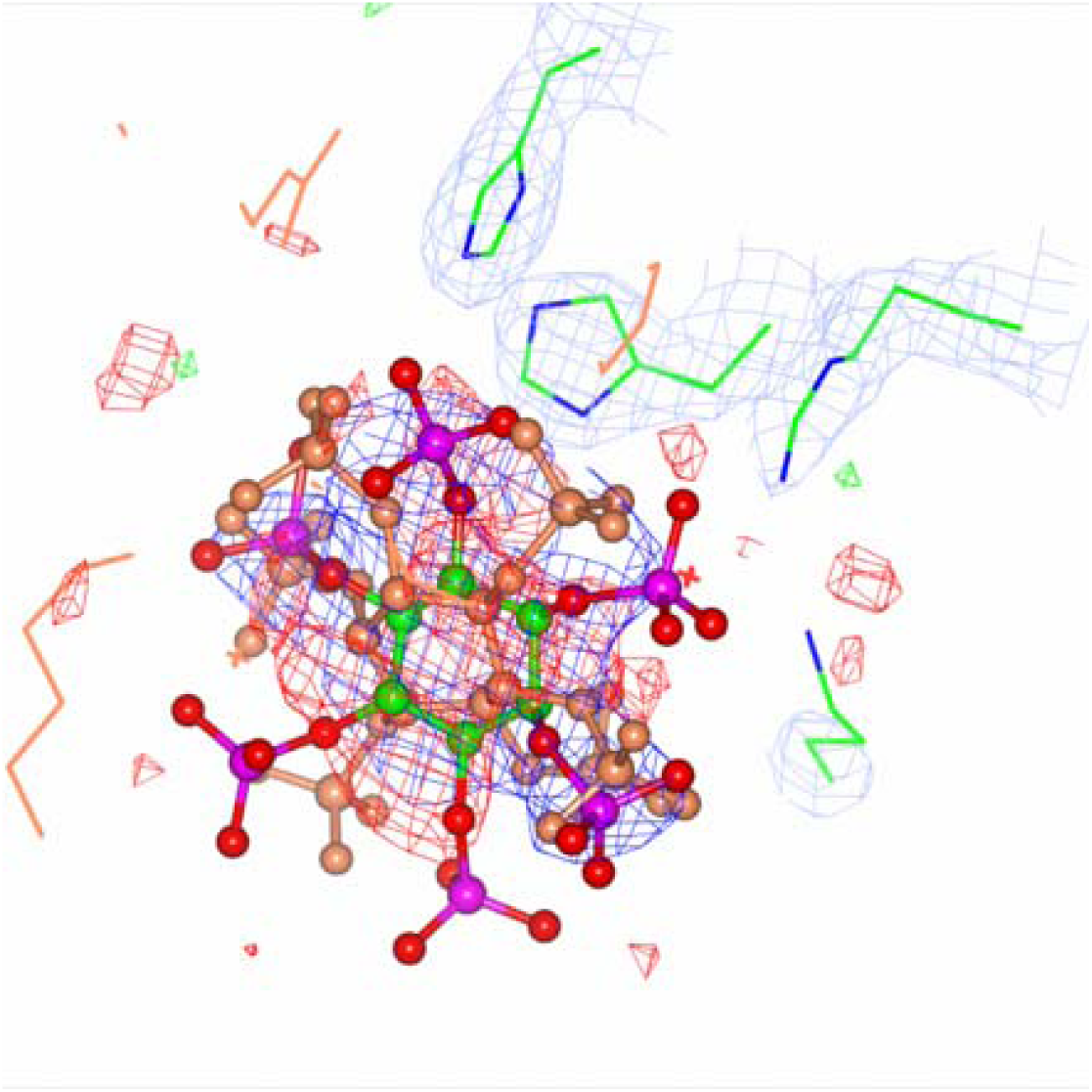
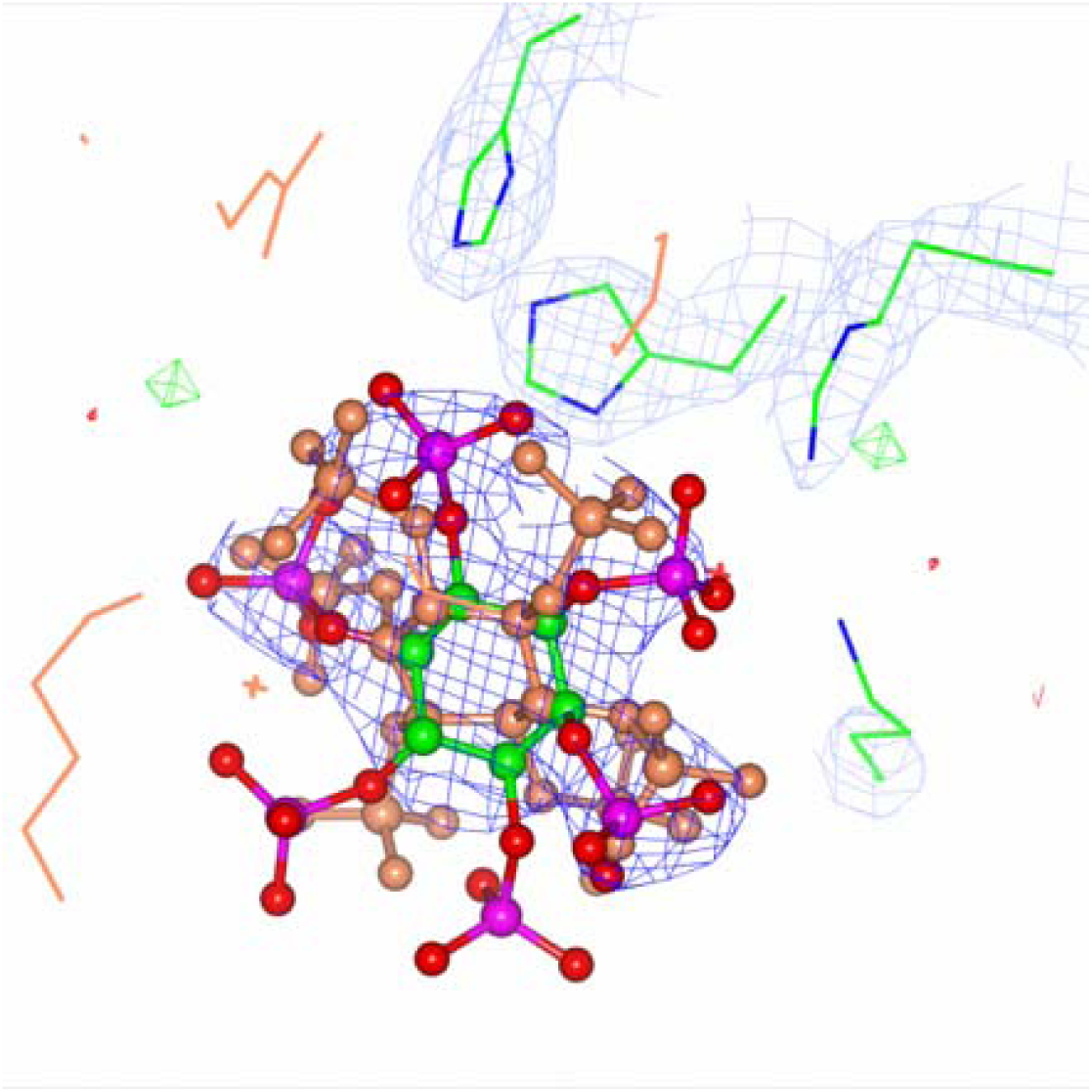
I6P ligand of 6B9B. (a) Fully occupied ligand with its symmetry mate. (b) half-occupied ligand with its symmetry mate.

#### Case 3: 6B9B

Occupancy of the ligand MAL of B chain of this PDB is estimated to be 0.414. Electron density shows there is no convincing evidence that fully occupied ligand exists in the crystal (Figure 9a). Refining the model without this ligand and adding waters according to the difference maps again cleaned up the density (Figure 9b).

**Figure 9.**
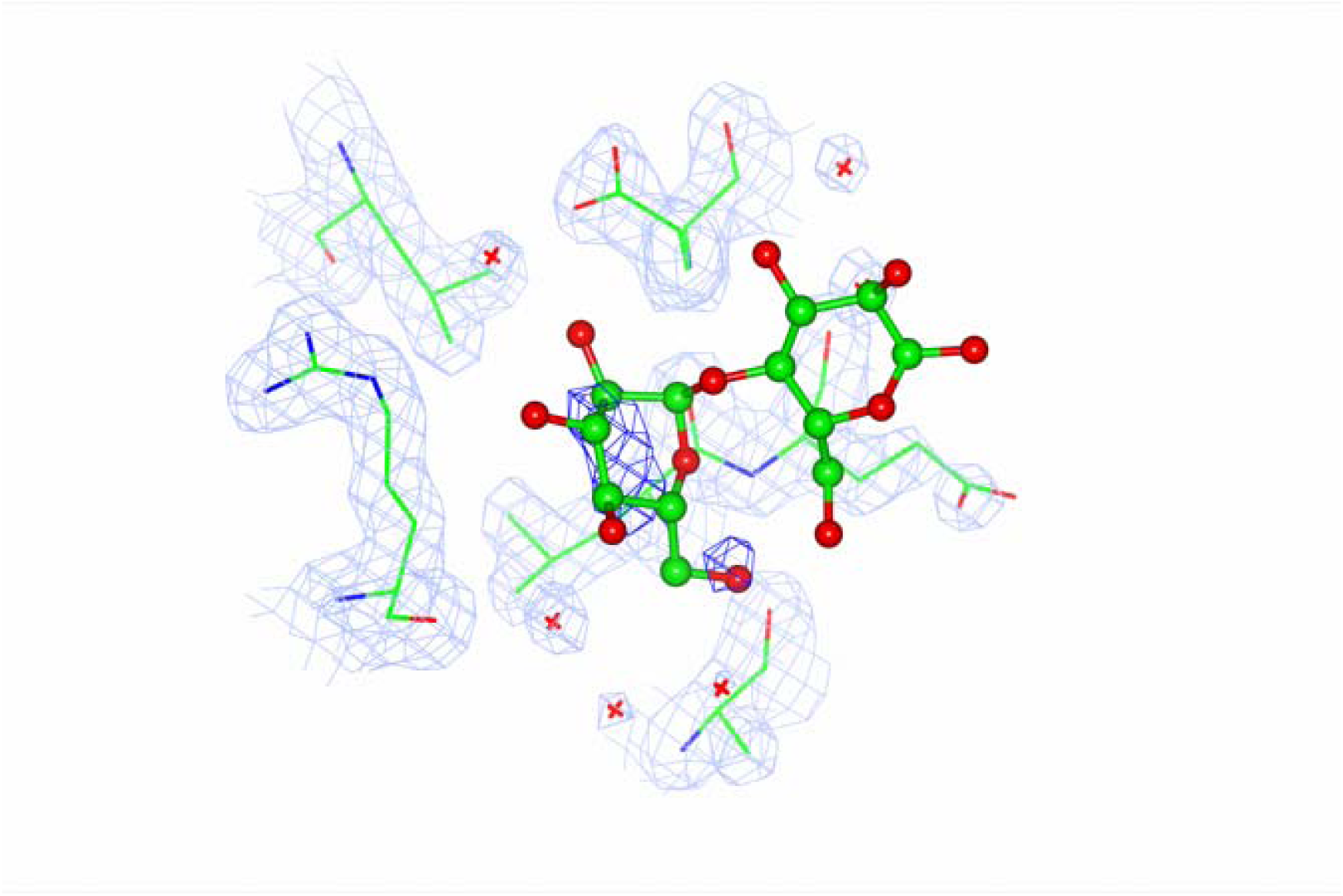
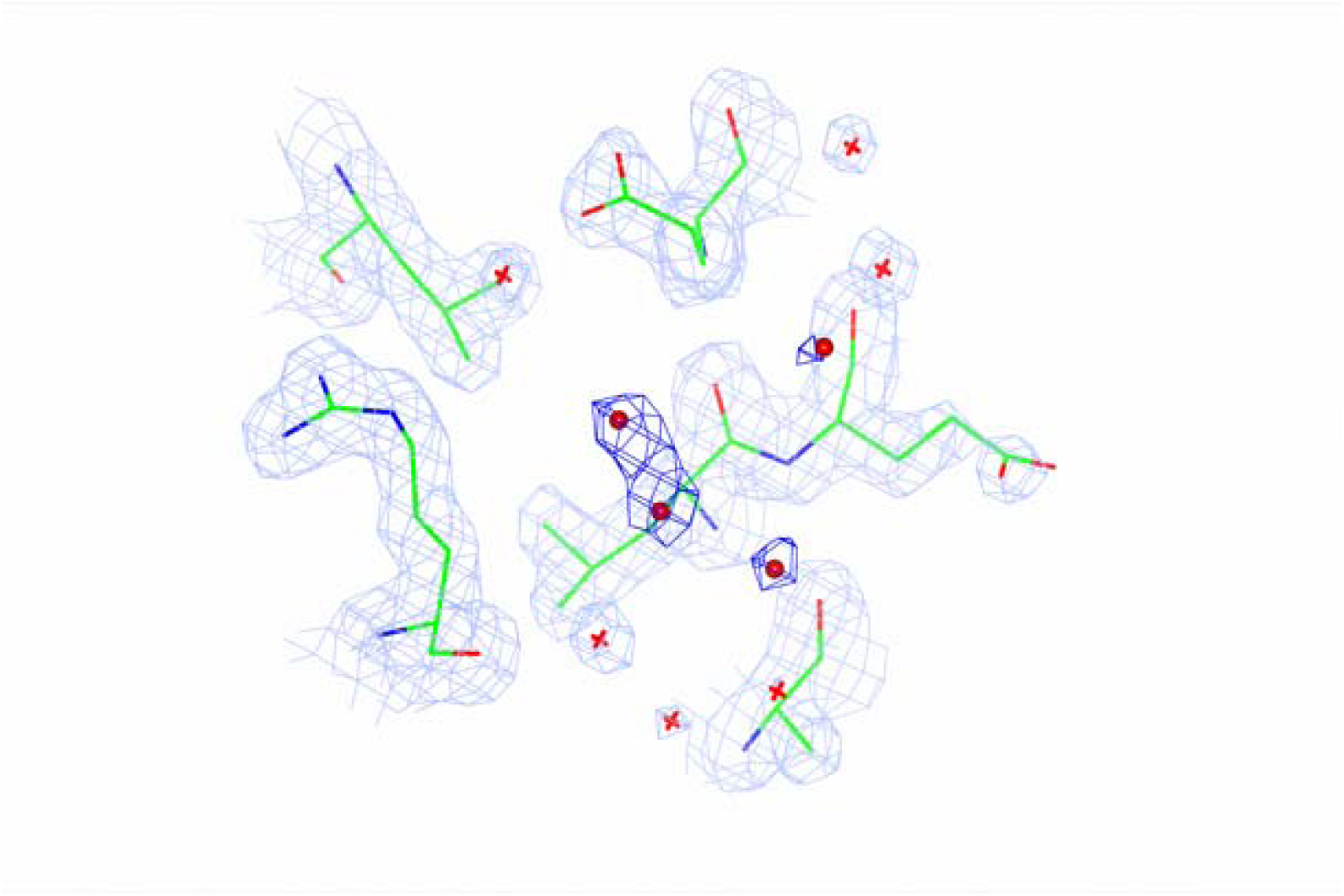
MAL ligand of 5ORJ. (a) The ligand before as it is in the PDB. (b) Water molecules placed using the difference map. These figures were produced using ccp4mg (McNicholas, 2011)

All these examples show that comparative analysis of ADPs of ligands with their neighbours can play a role in validation and has a potential in identifying wrong or disordered ligands.

## Conclusions

Many macromolecular structures from the PDB solved with X-ray crystallography exhibit multimodality of ADP distributions. The reasons for such behaviour are either wrongly modelled parts of the structure or different domains having different intermolecular contacts. In both cases parts of the molecule corresponding to the modes with large average ADPs should be inspected. Such ADP distributions are modelled using a mixture of SIGDs using Silverman method for the number of modes identification and expectation maximisation algorithm for the parameter estimation. ADPs of around 10% of the inspected PDB entries exhibit multimodality. Such multimodality also indicates that the resolution of maps corresponding to the different parts of the structure is different. In the limiting case, when atoms have been placed in wrong positions, resolution of the map is vanishing small. Analyses of the modes can shed some light into correctness, validity and mobility of different parts of the molecule, thus helping in validation and analyses of the resulting PDB structures. It can be expected that there will be more structures derived using cryoEM method exhibiting multimodality as variation of local resolution in these structures have been well documented (Kucukelbir et al, 2013).

The resolution and ADP dependent analysis of neighbouring atoms within structures has a potential in pinpointing mismodelled parts of the molecules. This tool can be used as a complementary validation during model building, refinement and deposition. Using this tool one can identify heavy atoms thus it can be used for modelling metal atoms. However, if this tool is used for identification of metal atoms then their coordination should also be accounted for.

Comparative analysis of ADPs of ligands and surrounding atoms using the algorithm developed in this work allows identification of potentially disordered and wrongly modelled ligands. Current approach uses the whole ligand as a unit. In practice there are many cases when only one part of a ligand is visible in the density. The algorithm can be extended to identify such cases by considering only local atom groups or local graph describing parts of the ligands. It should be emphasised that current algorithm assumes that the chemistry of ligands is correct. For general ligand validation one needs to use chemistry validation including stability of ligands, B values and density maps together to inform user about validity and nature of the ligands.

In future the use of the SIGD and local B value discrepancies in designing ADP restraints will be explored. It seems that using resolution dependent ADP restraints may result in better refinement. It could also be applied for atoms sufficiently far from each other.

Described algorithms have been implemented in the program *ToBvalid* which is available from *https://github.com/ToBvalid/* as an open-source software.

## Acknowledgements

This work was supported by MRC grant (MC_US_A025_0104) and Azerbaijan Academy of Sciences grant - decree № 5/9 dated on 15.03.2017. The authors also thank MRC-LMB, Cambridge, UK and IMBB of ANAS, Baku, Azerbaijan for creating encouraging research environment.

## Appendix A Estimation of the parameters of multimodal B value distributions

Let ***B*** be a vector of the sample of the data that comes from the population with the probability distribution as a mixture:

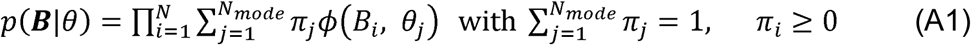

where *π*_*j*_ *s* are the mixture parameters and *θ*_*j*_ are parameters of *ϕ* (*B, θ*) corresponding to the mode *j, ϕ* (*B, θ*) is the parametrized family of distributions, *N* is the number of data points and *N*_*mode*_ is the number of modes. In the case of ADPs, the parameterised distribution is SIGD (2). Estimating parameters of (A1) directly is numerically unstable as extremely small and large values are summed together. To circumvent this problem an additional vector of a random variable – ***Z*** that is the belongness of each point to different modes is introduced. Then the resulting probability distribution of the augmented model is:

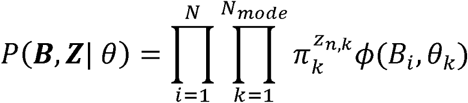

As a result, we have a product of the density of distribution which is easier to optimise, however new unknown parameters – ***Z*** has been introduced. This problem can now be solved using expectation and maximisation algorithm (Dempster et al, 1977; Bishop, 2006): estimate ***Z*** as expectation of the posterior probability distribution *P*(***Z*** | ***B, θ***) and estimate parameters ***θ*** using maximum likelihood estimation using the distribution P(*B* | ***Z, θ***) with fixed ***Z***. The algorithm for solution of this problem is well known (see for example Bishop, 2006). Here, we adapt this algorithm to estimate the parameters of the mixed SIGD:

1. Estimate the number and centroids of the modes using Silverman’s Test for Multimodality (Silverman, 1981) as implemented in *scipy*.
2. If *N*_*mode*_ > 1 then calculate peak heights using formula (3). It gives us Peak Height Distribution. Applying Expectation Maximisation algorithm with Gaussian Mixture Model (Bishop, 2006) estimate posterior distribution of belongness of each atom to each mode: *z*_*ij*_ *i* = 1..*N,j* = 1..*N*_*mode*_.
3. Using (*z*_*ij*_) apply EM algorithm for SIGD Mixture Model.
  a. Estimate the initial parameters of SIGD:
    i. Mixture parameters: 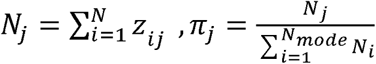
    ii. Find mean and minimum of B for each group: 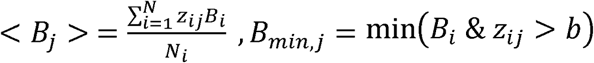, where *b* is a very small positive number
    iii. Set parameters: 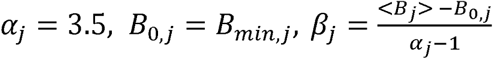
  b. Expectation step: 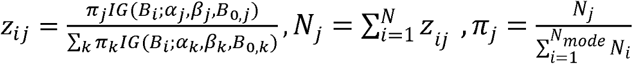
  c. Maximisation step: At this step we use only *z*_*ij*_ for which *B*_*i*_ > *B*_0,*j*_ otherwise corresponding *z*_*ij*_ set to 0. During summation to avoid negative arguments in the logarithms this fact is accounted for. For each mode calculate derivatives and do maximisation. Note that formulas are the same as in Masmaliyeva and Murshudov (2019) except they are now applied for each mode.
    i. The negative log-likelihood function has the form:

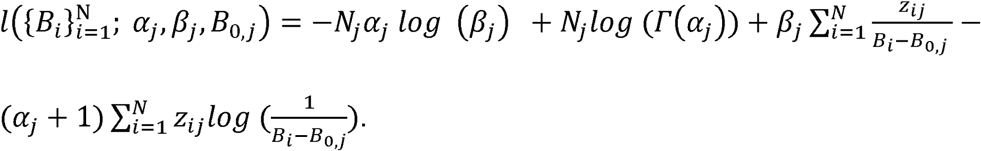
    ii. The first derivatives have the form:

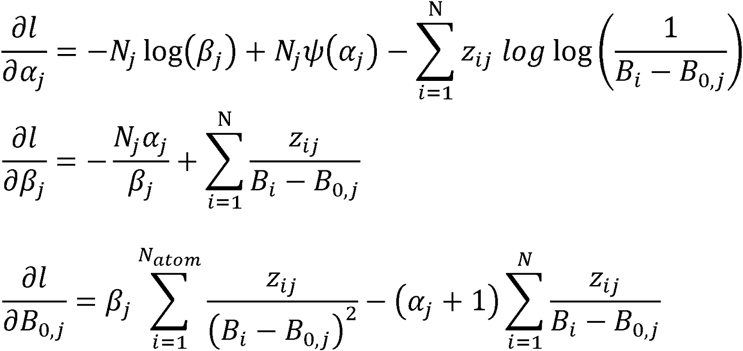
    iii. The expected Fisher information matrix has the form:

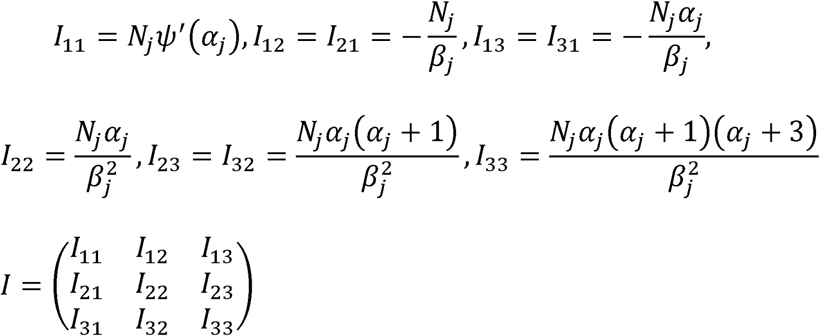
    iv. Find the shifts: *s* = −*I*^−1^*g*, where 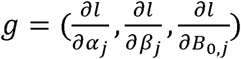 and apply them to the parameters *α*_*j*_, *β* _*j*_, *B*_0,*j*_)
    v. Repeat calculations until convergence
  d. Repeat b and c until convergence

## Appendix B. Local B value analysis

Estimation of occupancy of an atom in relation to its surroundings is done using total density difference (8) and simple statistics. The procedure consists of three steps:

1. The number and list of neighbours of each atom are calculated using *gemmi* library (Wojdyr, 2017); interatomic distance equal to 4.2Å is used as a default parameter. This can be adjusted by a user as an input parameter to the program *ToBvalid*.
2. If an atom has 3 or more neighbours, it is tested further. Values of relative peak height at the centre of atoms are calculated using the formula (8) for the atom in relation to neighbours. Let the corresponding relative occupancies of the atom be *c*_*0*_, *c*_*1*_ and *c*_*3*_ with respect to the median, first and third quartiles of ADPs of the neighbouring atoms. If *c*_*0*_ > 1.2 and *c*_*1*_ > 1.01 then atom is considered heavier than neighbours, if *c*_*0*_< 0.8 and *c*_*3*_ < 0.99 then this atom is considered lighter than neighbours. In both cases optimal occupancy - *c*_*0*_ is reported. The parameters are default values that have been selected by trial and error.
3. If the inspected atom is oxygen of a water molecule and it has six or more neighbours then it is marked as an atom with unusual behaviour atom. These are candidates for metals.

## Notes

### Competing Interest Statement

The authors have declared no competing interest.

